# Individual cell-based modeling of tumor cell plasticity-induced immune escape after CAR-T therapy

**DOI:** 10.1101/2020.06.20.163170

**Authors:** Can Zhang, Changrong Shao, Xiaopei Jiao, Yue Bai, Miao Li, Hanping Shi, Jinzhi Lei, Xiaosong Zhong

## Abstract

Chimeric antigen receptor (CAR) therapy targeting CD19 is an effective treatment for refractory B cell malignancies, especially B cell acute lymphoblastic leukemia (B-ALL). The majority of patients achieve a complete response following a single infusion of CD19-targeted CAR-modified T cells (CAR-19 T cells); however, many patients suffer relapse after therapy, and the underlying mechanism remains unclear. To better understand the mechanism of tumor relapse, we developed an individual cell based computational model for tumor cell plasticity and the heterogeneous responses to the CAR-T treatment. Model simulations reproduced the process of tumor relapse, and predicted that CAR-T stress-induced cell plasticity can lead to tumor relapse in B-ALL. Model predictions were verified by applying the second-generation CAR-T cells to mice injected with NALM-6-GL leukemic cells, in which 60% of the mice relapse within 3 months, and relapsed tumors retained CD19 expression but exhibited a subpopulation of cells with CD34 transcription. These findings lead to a mechanism of tumor replace by which CAR-T treatment induced tumor cells to transition to hematopoietic stem-like cells (HSLCs) and myeloid-like cells and hence escape of CAR-T targeting. The computational model framework was successfully developed to recapitulate the individual evolutionary dynamics, which could predict clinical survival outcomes in B-ALL patients after CAR-T therapy.

## Introduction

Cancer immunotherapy based on genetically engineered T cells has been used successfully to treat refractory B cell malignancies, especially acute lymphoblastic leukemia (ALL)(1–3). In this strategy, the T cell genome is modified by the integration of viral vectors or transposons encoding chimeric antigen receptors (CARs) that direct tumor cell killing(4,5). The application of CAR-T therapy is limited by two major challenges: cytokine release syndrome (CRS), which is potentially life-threatening for patients and hence limits the number of CAR-T cells that can be transferred, and disease relapse, which occurs for various reasons(6–11). A detailed understanding of the disease relapse process is certainly important for the improvement of CAR-T cell therapy.

CAR-T therapy targeting CD19 has been proved to be an effective therapy for B cell acute lymphoblastic leukemia (B-ALL). Following CD19 CAR-T treatment, complete remission is often observed within 3 months; however, a significant number of patients relapse in long-term follow-up(2,12–26). These disease recurrences are associated with various phenomena, including mutation or alternative splicing of CD19, or the presence of CD19^-^ cancer cells(27–29), or CD19^+^ relapse(2,13,14,19,25). Some relapsed patients show myeloid leukemia switches(30–32). In a study of human non-Hodgkin’s lymphoma (NHL) treated with CAR-19 CD8^+^ T cell (CTL) therapy, the acquisition of resistance was independent of the downregulation/loss of CD19 expression and was presumably due to deregulated apoptotic machinery in anti-CD19 CAR CTL-resistant NHL sublines (R-NHL)(33). The differentiation plasticity of hematopoietic cells has been increasingly recognized in recent years(34,35), and some patients develop mixed-phenotype acute leukemia(36–38), which shows aberrant expression of antigens from different lineages. Dual CARs, such as CD22-targeted or dual CD19- and CD123-targeted T cells, are often applied in relapsed patients(39,40). The reasons for tumor relapse, especially CD19^+^ relapse, are not fully understood, and detailed studies of the dynamic process after CAR-T cell infusion are necessary to uncover the underlying mechanisms of immune escape following CAR-T therapy.

Drug treatments can promote cellular plasticity, which may help cancer cells evade detection and treatment(41). Lineage switches after CAR-T therapy in B-ALL shown a possible response of CAR-T stress-induced tumor cell plasticity. We asked how CAR-T stress-induced cell plasticity may result in the immune escapes after CAR-T therapy? How the process of tumor relapse may depend on the tumor cell plasticity and heterogeneous responses to CAR-T stress? How can we perform personal prediction of the outcome of CAR-T therapy? Experimentally, it is difficult to track details of cell plasticity during tumor relapse. Here, to better understand the mechanism of cell plasticity-induced tumor relapse, we developed an individual cell based computational model for the heterogeneous responses of tumor cells to the CAR-T treatment based on a hypothesis of CAT-T treatment induced tumor cell plasticity and immune escape. Model simulations reproduced the process of tumor relapse, and shown that CAR-T stress-induced cell plasticity can lead to tumor relapse. Phenotype changes in relapsed tumor cells were further verified by mouse experiments, which shown the existence of CD34^+^ and CD123^+^ cells in tumor cells 72 days after CAR-T cells infusion. Based on model simulations, tumor relapse can be predicted by dynamic parameters associated with individual responses to CAR-T treatment.

## Materials and Methods

### CAR-T treatment in mice injected with NALM-6-GL leukemic cells

We developed a mouse model and applied the modified second-generation CD19-specified CAR-T cells CD19-28z(42–44). The functions of the modified CD19-28z T cells in response to tumor cells were tested through *in vitro* experiments. When CD19-28z T cells were co-cultured with K562-CD19 or NALM-6-GFP-Luc cells (NALM-6-GL), significantly high CD107a expression was observed in both CD8+ and CD4+ CAR-T cells, and IFN-γ was specifically produced (**Figure S1**). Hence, the CD19-28z T cells were effectively activated by specific antigen CD19 cell lines and primary ALL cells.

We injected mice with NALM-6-GL tumor cells on day 1, followed by CD19-28z CAR-T cells infusion on days 2, 3, 4, and 12. All mice showed complete remission (CR) in the first few weeks, but 60% (16/27) showed tumor relapse in long-term tracking. The tumor burden was assessed as previously described through bioluminescence imaging using a Xenogen IVIS Imaging System (Xenogen) with Living Image software (Xenogen) (45). Both dorsal and ventral views were obtained for all animals. To examine the phenotype of the tumor cells in the relapse mice, GFP+ tumor cells isolated from the bone marrow were sorted by flow cytometry.

Detail experimental protocols were given in the Supplementary Text.

### Individual cell-based modeling of tumor relapse

We developed an individual cell based computational model for the heterogeneous tumor cell responses to CAR-T treatment. The model simulates the biological process of tumor growth as a multiple cells system, in which each cell is undergoing proliferation, death, or terminal differentiation with their own rates depending on expression levels of the marker genes in the cell as well as the CAR-T signaling.

The model considers all tumor cells in the bone marrow, which are divided into three components: hematopoietic stem-like cells (HSLCs), progenitors, and terminally differentiated cells (TDCs) (**Figure 1a**). Different type cells differ in their abilities for self-renewal and terminal differentiation. HSLCs are quiescent cells with low self-renewal ability, the highest potential to produce diverse progeny, and no terminal differentiation; progenitor cells are highly self-renewable, and can differentiate into TDCs; and TDCs have lost the capability of self-renewal, circulate through the body, and are prone to cell death. Upon CAR-T treatment, we assumed that tumor cells are killed by CAR-T cells with a death rate depending on the CAR-T signaling strength of each individual cell. The cell-to-cell variance in CAR-T signaling may be dependent on the expression of surface markers on target cells (**Figure 1b**), which vary over cell regenerations.

**Figure 1.**
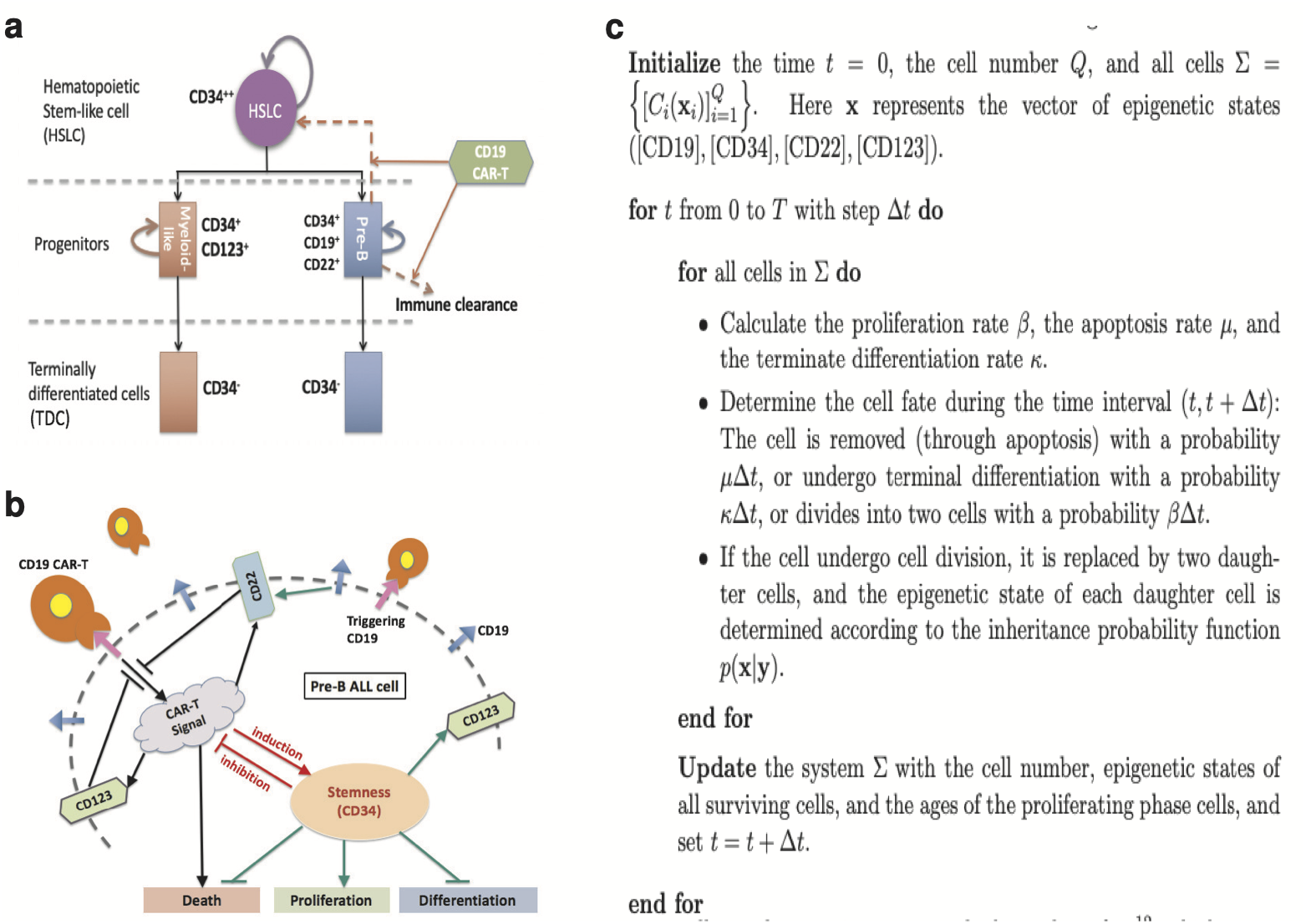
Schematic representation of the single-cell based computational model. **a.** Main hypotheses of the computational model (see the Supplementary Text S2). All tumor cells were classified as hematopoietic stem-like cells, progenitors, or terminally differentiated cells. CD19-28z CAR-T cells can kill CD19^+^ pre-B tumor cells and induce the transition from pre-B cell to hematopoietic stemlike cell by promoting the expression of CD34. **b.** Major assumptions of the interactions between the marker genes, CAR-T cell signaling, and cell behavior. Green arrows show the interactions independent of the CAR-T signal, black and red arrows represent the interactions associated with the CD19-28z CAR-T. The two key assumptions, the induction of stemness by the CAR-T signal and immune escape of the stem-like cells, are marked red. **c.** Numerical scheme of model simulation.

In the model, we introduced key assumptions that CAR-T stress-induced tumor cells to transition to HSLCs (by promoting CD34 expression) and myeloid-like cells (by promoting CD123 expression) and hence escape of CAR-T targeting. Heterogeneity of each cell was represented by the relative expression levels of marker genes CD19, CD22, CD34, and CD123, which play important roles in the CD19 CAR-T cell response and cell lineage. The proliferation rate *β* and the differentiation rate *κ* depend on CD34 expression level through

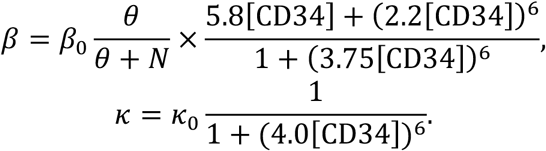

Here *N* means the total tumor cell number. CAR-T signals are dependent on the expression of these marker genes

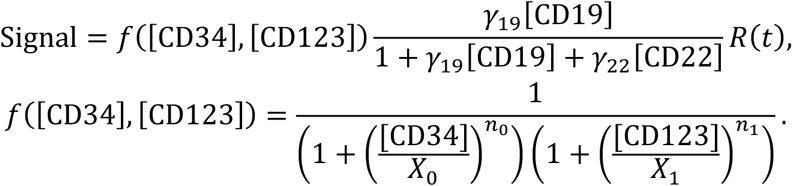

Here *R*(*t*) is the predefined CAR-T activity (referred to Supplementary Text, equation (1)). CD34 and CD123 are marker genes of stem-like cells and myeloid-like cells, respectively, which were assumed to inhibit the CAR-T signaling. The apoptosis rate *μ* includes a basal rate *μ*_0_ and a rate associated with the CAR-T signal

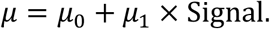

In model simulations, we omitted the complex yet incomplete signaling pathways for the marker gene expressions, and introduced phenomenological descriptions for the random transitions of gene expressions. We assumed that the expression of the marker genes randomly changed during cell cycling following a transition probability of Beta-distributions, of which the shape parameters were dependent on both the state of mother cells and the CAR-T signaling. For example, we assumed that both CD19 and the CAR-T signal promote CD34 expression, so that given the expression of CD34 (*u_k_*) at the *k*^th^ cycle, the average expression level at the (*k* + 1)^th^ cycle (denoted by *u*_*k*+1_) is

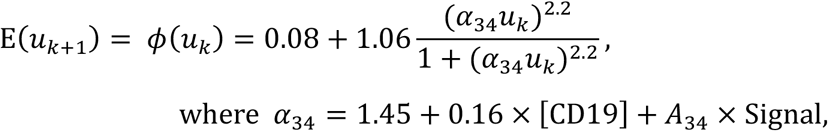

and the variance is

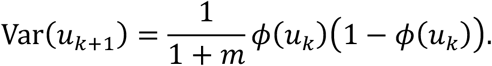

Let the shape parameters

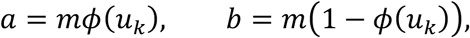

the expression level *u*_*k*+1_ is a Beta-distribution random number with probability density function

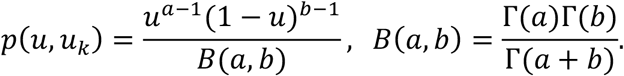

Formulations for the other genes were given similarly (Supplementary Text).

Model parameters were adjusted in accordance with the experimentally observed tumor relapse dynamics and tumor cells marker gene expression through flow cytometry analysis (**Table S3** in the Supplementary Text).

To reproduce the tumor relapse process, we performed model simulations to mimic the process of tumor cells proliferation, death, and terminally differentiation, and cell plasticity during cell proliferation (**Figure 1c**). For detail formulations, parameter estimations, and numerical schemes of the model, referred to the Supplementary Text.

## RESULTS

### Tumor relapse after CD19-targeted CAR-T treatment in mice injected with NALM-6-GL leukemic cells

We developed a mouse model and applied second-generation CAR-T cells to mice injected with NALM-6-GL leukemic cells. The functions of the modified CD19-28z T cells in response to tumor cells were tested through *in vitro* experiments. CD19-28z T cells were co-cultured with various types of target cells at different effector to target ratios (E:T ratios). The target cells were effectively killed when the E:T ratio was larger than 5:1 (**Figure S1c**), which confirmed the CD19-targeted lysis of the CAR-T cells. Moreover, the carboxyfluorescein succinimidyl ester (CFSE) proliferation assay showed that the CD19-28z T cells underwent multiple rounds of cell division when co-cultured with NALM-6-GL, but not with K562 (**Figure S1d**). This result revealed the proliferative ability of CD19-28z T cells in response to cognate-antigen recognition on target cells.

To investigate CD19-28z CAR-T treatment to B-ALL, we applied a rapid expansion protocol (REP) (46,47) to obtain clinically relevant numbers of T cells for adoptive T cell transfer and applied NALM-6-GL to the mouse model. We injected mice with NALM-6-GL tumor cells on day 1, followed by T cell infusion on days 2, 3, 4, and 12 (**Figure 2a**). The mice were divided into two groups: infusion with control NGFR-28z T cells and infusion with CD19-28z T cells. In the NGFR-28z groups, all mice showed similar dynamics of rapid tumor growth after injection and died within 7 weeks (**Figure 2b-c**). The mice treated with CD19-28z T cells, however, showed diverse responses in the tumor cell population dynamics. All mice showed complete remission (CR) in the first few weeks, but 60% (16/27) showed tumor relapse in long-term tracking (**Figure 2b-c**).

**Figure 2.**
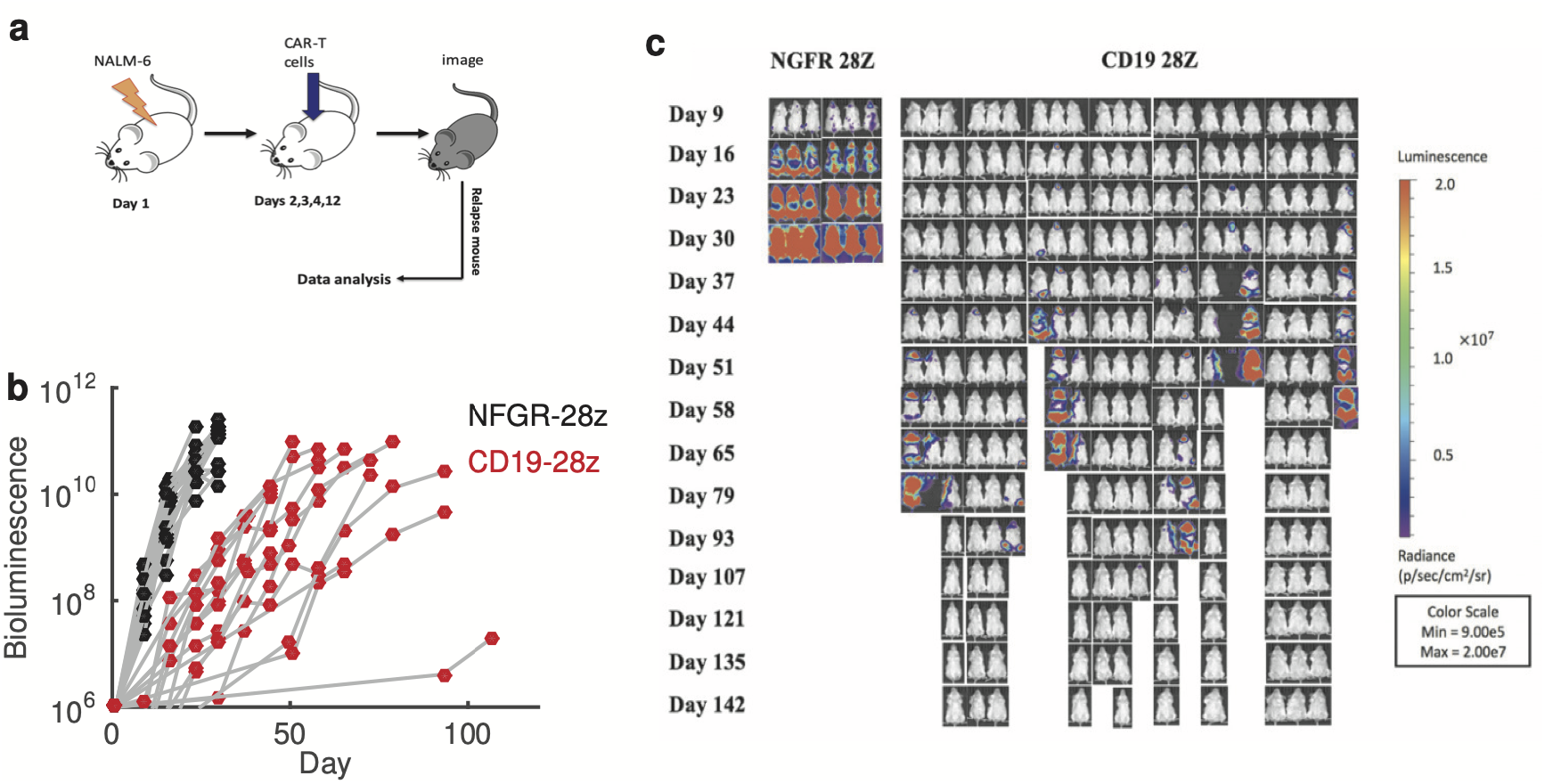
Tumor relapse in mice treated with CAR-T cells. **a.** Experimental process. On day 1, 1×10^6^ NALM-6-GL cells were injected through the tail vein into the NOD-SID mice. On days 2,3,4, and 12, 5×10^6^ CD19-28z CAR-T cells were injected through the tail vein into each NALM-6-GL-bearing NOD-SCID mouse. Cancer development was measured with bioluminescence imaging, and tumor cells were isolated from the bone marrow for further flow cytometry analysis. **b.** Bioluminescence imaging of mice to detect tumor progresses. **c.** Evolution of tumor progresses represented by bioluminescence data. The data were obtained from mice receiving control NGFR-28z T cells (black) and mice that relapsed after CD19-28z CAR-T cell treatment (red).

To investigate the phenotype of relapsed tumor cells, we isolated the GFP^+^ tumor cells from the bone marrow of relapsed mice at days 37 and 72, respectively, and sored the isolated cells by flow cytometry. Most cells from both the early (day 37) and late (day 72) stages showed CD19 positivity (**Figure 3a**), as confirmed by CD19 immunohistochemistry (**Figure 3b**). Flow cytometry showed the existence of CD19^+^CD34^+^ and CD123^+^CD34^+^ tumor cells in the CD19-28z-treated relapsed mice, which were not presented in NGFR-28z-treated mice (**Figure 3c-f**). These results shown the presence of CD34^+^ tumor cells in relapsed mice, which suggest the possibility of tumor cell plasticity after CAR-T treatment.

**Figure 3.**
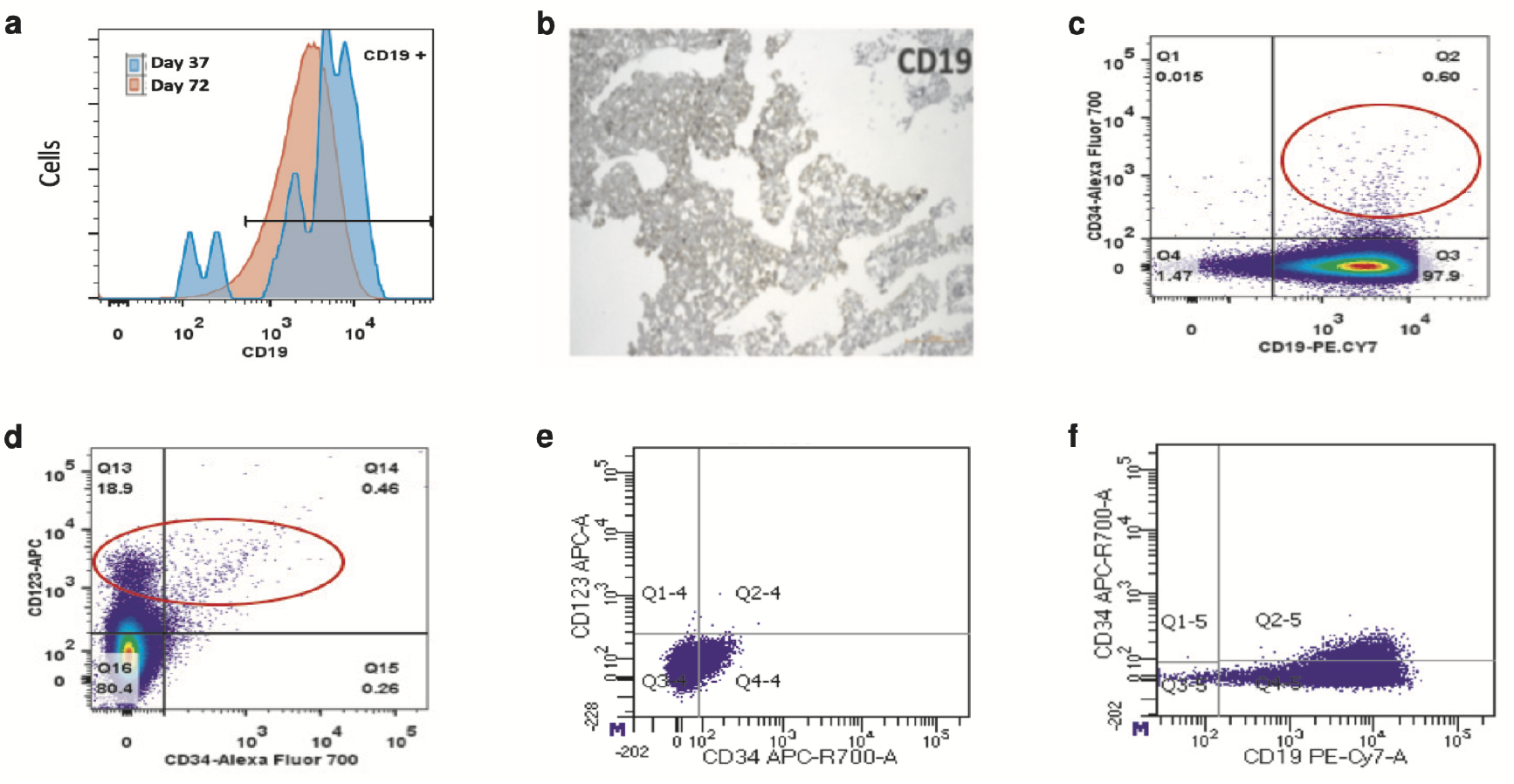
Phenotype of relapsed tumor cells. **a.** Flow cytometry analysis of CD19 expression on the tumor cells pooled from three relapsed mice at days 37 and 72, respectively. **b.** CD19 immunohistochemistry of bone marrow sections obtained from CD19-28z CAR T cell-treated mouse (week 4). **c.** Flow cytometry plot of CD34 expression versus CD19 expression in the bone marrow tumor cells isolated from the relapsed mice at day 72. Red circle indicates the cells with phenotype CD34^+^CD19^+^. **d.** Flow cytometry plot of CD123 expression versus CD34 expression in the bone marrow tumor cells isolated from the relapsed mice at day 72. Red circles indicate the cells with phenotype CD123^+^CD34^+^ and CD123^+^CD34^-^. **e.** Flow cytometry plot of CD123 expression versus CD34 expression in the bone marrow tumor cells isolated from the NGFR-28z-treated mice at day 30. **f.** Flow cytometry plot of CD34 expression versus CD19 expression in the bone marrow tumor cells isolated from the NGFR-28z-treated mice at day 30.

### Model simulations of tumor relapse after CAR-T treatment

In the proposed individual cell-based model, we introduced key assumptions of CAR-T stress-induced cell plasticity, CAR-T signals promote the expressions of CD34 and CD123, so that tumor cells can escape CAR-T targeting. To verify the proposed hypothesis, we mimicked the experimental process to simulate tumor growth after NALM-6-GL cells injection and CAR-T treatment. In simulations, we initialized the system with 1×10^6^ tumor cells, which were all CD19^+^ with low expressions of CD34, CD22, and CD123. First, we turned OFF the CAR-T signaling and adjusted the parameters for cell proliferation and death rates to fit the experimental data from NGFR-28z-treated mice (**Figure S2**). Next, we turned ON the CAR-T signaling and adjusted the parameters related to tumor cells response to fit the process of tumor relapse (**Figure 4a**). Simulations reproduce the diverse tumor cell population dynamics of different relapsed mice after CD19-28z T cell infusion, which revealed heterogeneous responses among different mice. In simulations, the heterogeneous responses in different mice were represented by varying the two parameters, the ability of CAR-T stress-induced CD34 expression (A_34_) and the inhibition of CAR-T signals by CD34 expression (*X*_0_) (**Figure 1b**). Here, the parameter for CAR-T stress-induced CD34 expression measures the stem-like cell plasticity induced by CAR-T stress, and the inhibition of CAR-T signaling by CD34 expression indicates the capacity for immune escape of stem-like cells. In additional to the relapse process, the individual cell-based model enables us to simulate the expression levels of marker genes at single cell level, which are comparable with experimental data obtained from the flow cytometry analysis (**Figure 4b**).

**Figure 4.**
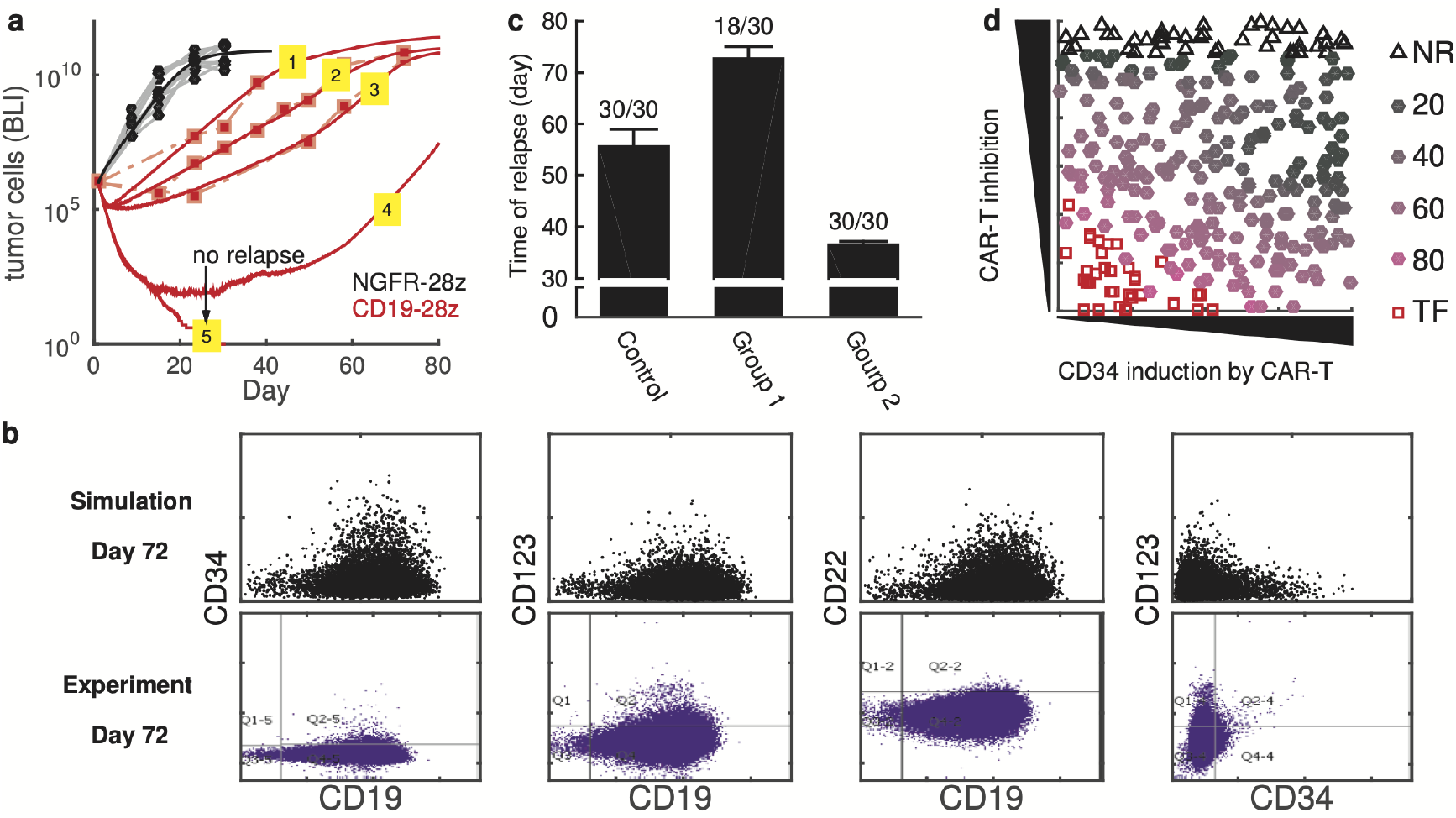
Simulation results and experimental data for a mouse with the CD19-28z CAR-T treatment. **a.** Tumor growth after the NALM-6-GL injection and CD19-28z CAR-T treatment. The markers show the bioluminescence imaging (BLI) experimental data, and the solid curves were obtained from the model simulation. Simulated tumor cell numbers are scaled (x10^6^) to compare with the BLI data. Black indicates mice treated with the control NGFR-28z, and red indicates mice treated with CD19-28z. Here, simulation results for 5 mice treated with CD19-28z are shown; #1-#3 were fit to the data from three mice, #4 models the case with slow relapse, and #5 models the case of tumor cell clearance. Refer to **Table S4** in the Supplementary Text S2 for the parameters used for #1-#5. **b**. Scatter plots marker gene expression at day 72, from the simulation (upper panel) and the flow cytometry analysis (bottom panel). Data obtained from the mice #2 in a. **c.** Time of relapse under the three conditions: Control (*A_34_=0.4, X_0_=0.4*(*±20%*)), Group 1 (*A_34_=0, X_0_=0.4*(*±20%*)), and Group 2 (*A_34_=0.4, X_0_=0.2*(*±20%*)). In each case, we performed 30 independent runs, each of which the parameter *X_0_* for the inhibition the CAR-T signal by CD34 varied over a given range. Numbers on the error bar show the cases with tumor relapse in the 30 runs. **d.** Simulated relapse time with different parameters for CD34 induction (*A_34_*) by the CAR-T and CAR-T signal inhibition by the stem-like cells (*X_0_*). Black triangles show the cases with no remission (NR) after the CAR-T cell injection, red squares show the cases became tumor free (TF) after CAR-T treatment, and solid circles show the cases with remission at the beginning and tumor relapse in the later stages. Here, increasing the CD34 induction by the CAR-T cells corresponds to an increasing of *A_34_*, and increasing the CAR-T signal inhibition corresponds to a decreasing of *X_0_*. The color of the solid circles indicates the day of relapse. The time of relapse was defined as the time when the tumor cell number regain the initial value after early remission.

We further changed the values of *A*_34_ and *X*_0_ to investigate how the dynamics of tumor relapse depends on these two parameters. When we turned the CD34 induction off (*A*_34_ = 0), the simulated relapse rate significantly decreased; and in the relapsed cases, the time to relapse was clearly postponed (**Figure 4c**, Group 1 vs. Control). By contrast, a decrease in the parameter *X*_0_ resulted in faster relapse (**Figure 4c**, Group 2 vs. Control). When we varied both parameters, an outcome of complete tumor cells clearance can be achieved when *A*_34_ was small and *X_0_* was large enough, which represent the conditions of weakenness the CD34 induction by CAR-T and the CAR-T signal inhibition of stem-like cells (**Figure 4d**). Moreover, model simulations predicted the situations of no remission when *X*_0_ was small so that CAR-T signal inhibition by CD34 was strong enough (**Figure 4d**). These results shown that the diverse responses of different mice to CAR-T treatment can be explained by the variance in the interactions between the CAR-T signal and tumor cell stemness (**Figure 1b**).

### The *in-silico* process of tumor relapse

Our model provides an *in-silico* laboratory to explore the process of tumor relapse. Based on the simulated cell population dynamics (**Figure 4a**), a typical process of tumor relapse without CD19 loss includes three stages, beginning with CR at the early stage, followed by a critical transition to the relapse phase, and finally an accelerated growth after CAR-T cell exhaustion (**Figure S3**). During tumor relapse, most tumor cells maintained a high level of CD19 expressions, and the CD34 distribution showed obvious switches from low to high levels in the early stage and back to the lower level distribution in the later stage after CAR-T cell exhaustion (**Figure 5a**).

**Figure 5.**
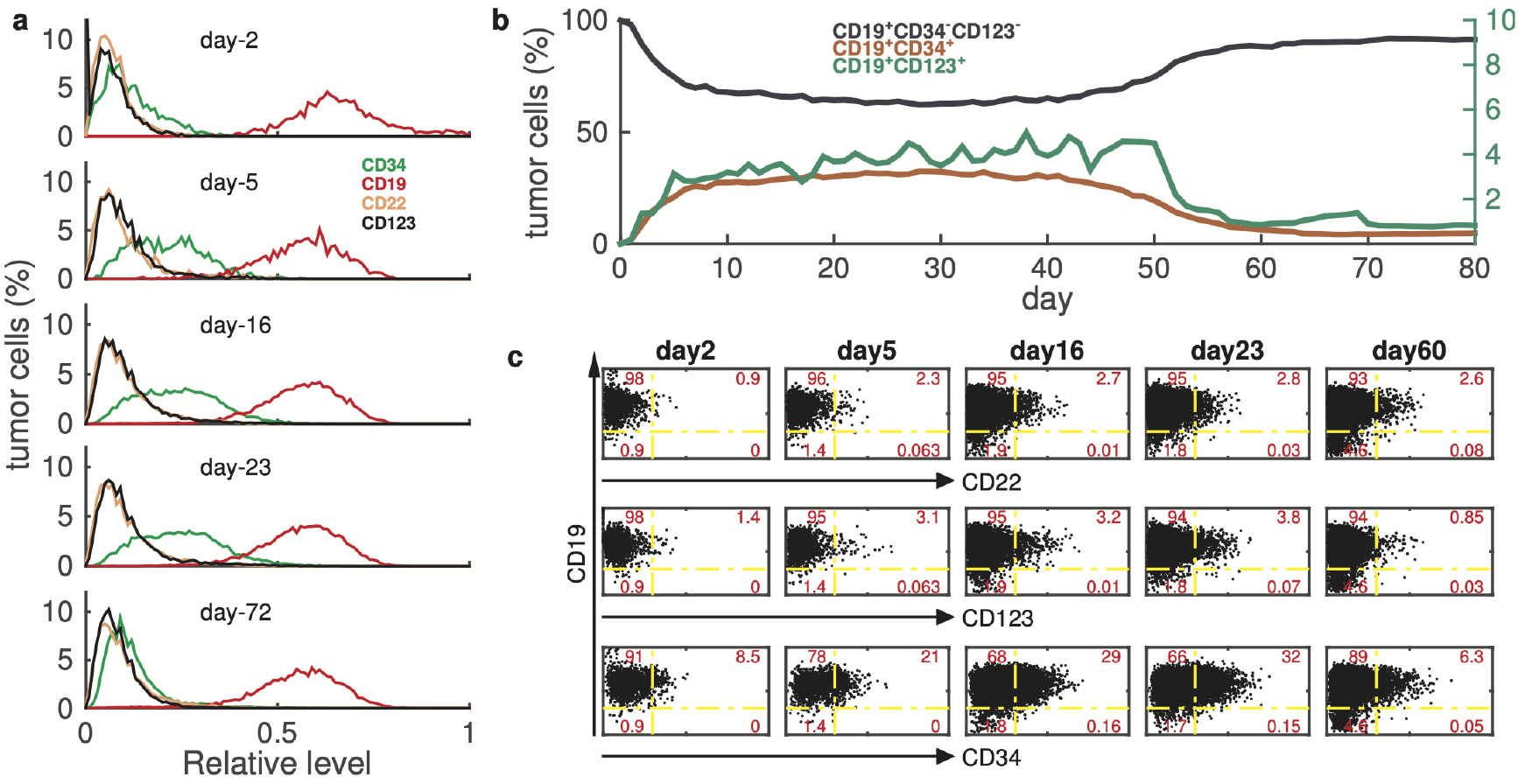
*in silico* process of tumor relapse. **a.** Simulated distribution of the marker genes CD34, CD19, CD22, and CD123 at days 2, 5, 16, 23, and 72 post-tumor cell and CAR-T cell injections. **b.** Time evolution of the percentages of the three subpopulations of cells: original tumor cells (CD19^+^CD34^-^CD123^-^), hematopoietic stem-like cells (CD19^+^CD34^+^), and myeloid-like cells (CD19^+^CD123^+^). **c.** Simulated relative levels of CD22, CD123, CD34, and CD19 in tumor cells during the tumor relapse process. Numbers show the percentages of cells in each quadrant. Data obtained from the mice #2 in **Figure 4a**.

We further analyzed the dynamics of various subpopulation cells during tumor relapse (**Figure 5b**). After the CAR-T infusion, the CD19^+^CD34^-^CD123^-^ cell number decreased immediately due to CAR-T lysis, while the CD19^+^CD34^+^ and CD19^+^CD123^+^ cells significantly increased during the relapse phase. At the latter stage, the CD19^+^CD34^+^ and CD19^+^CD123^+^ subpopulations decreased, whereas CD19^+^CD34^-^CD123^-^ cell number obviously increased, regaining the original phenotype of the injected NALM-6-GL cells. These results revealed the dynamics of cell lineage switches during tumor relapse, with lymphoid pre-B cells (CD19^+^CD34^-^CD123^-^) switch to a stem-like phenotype (CD19^+^CD34^+^) after the CAR-T treatment, followed by further switch to a myeloid-like cell phenotype (CD34+CD123^+^). The stem-like cells regenerated lymphoid pre-B cells after the CAR-T exhaustion/disappearance.

To explore cell plasticity during tumor relapse, we analyzed the relative expressions of marker genes at single cell level along the simulated relapse process. From the simulated process (mouse #2 in **Figure 4a**), the antigen CD19 maintained high level expression in most tumor cells, the fraction of CD34+ subpopulation cells obviously increased from days 5 to 23 after CAR-T cells infusion (**Figure 5c**), in agree with the above analysis. Moreover, there were small subpopulations of cells that showed upregulation of CD22 and CD123 expressions on days 5-23 after the CAR-T cells infusion (**Figure 5c**); but the changes in the CD123 and CD22 expression distributions were imperceptible over the relapse process (**Figure 5a**). These results show a process of tumor cell plasticity with upregulation of CD34 expression in response to CAR-T stress.

### Predictability of the computational model

In the above simulations, tumor relapse dynamics in different mice can be fitted by varying the parameters *A*_34_ and *X_0_* for the heterogenous responses in individual mice. This raise a possibility of predicting the outcome of CAR-T treatment by identifying the personalized parameters through an estimation of parameter values based on a short-term observation after CAR-T infusion(48). Clinically, it is crucial to predict, according to the responses at early stage after CAR-T infusion, whether the patient would be cured with tumor cells free, or, if otherwise, the day of tumor relapse.

To test the predictability based on *in silico* relapse process, we examine how the days of tumor relapses may depend on the heterogeneous response at different mice. We varied the parameters *A*_34_ and *X_0_* to mimic different mice (0 < *A*_34_ < 0.8,0 < *X_0_* < 0.6), and analyzed the cases showed initially remission with tumor cells decreasing (**Figure 6a**). For each simulated case, we calculated the relative tumor burden *(T)* and the fraction of CD34^+^ tumor cells (*f*_CD34_) at day 5 after CAR-T infusion (dashed line in **Figure 6a**). Here *T* measures the relative tumor cells with respect to the initial number before CAR-T infusion, and *f*_CD34_ is the fraction of tumor cells with CD34>0.3. **Figure 6b** shows *f*_CD34_ and *T* for each simulated case, the fraction *f*_CD34_ shows nonlinear correlation with the tumor burden *T* when *T* is small (*T* < 10^-2^). Moreover, the cases developed to tumor cells free usually have low relative tumor burden (*T* < 10^-2^) and small fraction of CD34^+^ cells. We further calculated the timing of tumor cells clearance, which show a Poisson distribution with a mean of about 30 days after CAR-T cells infusion (**Figure 6c**).

**Figure 6.**
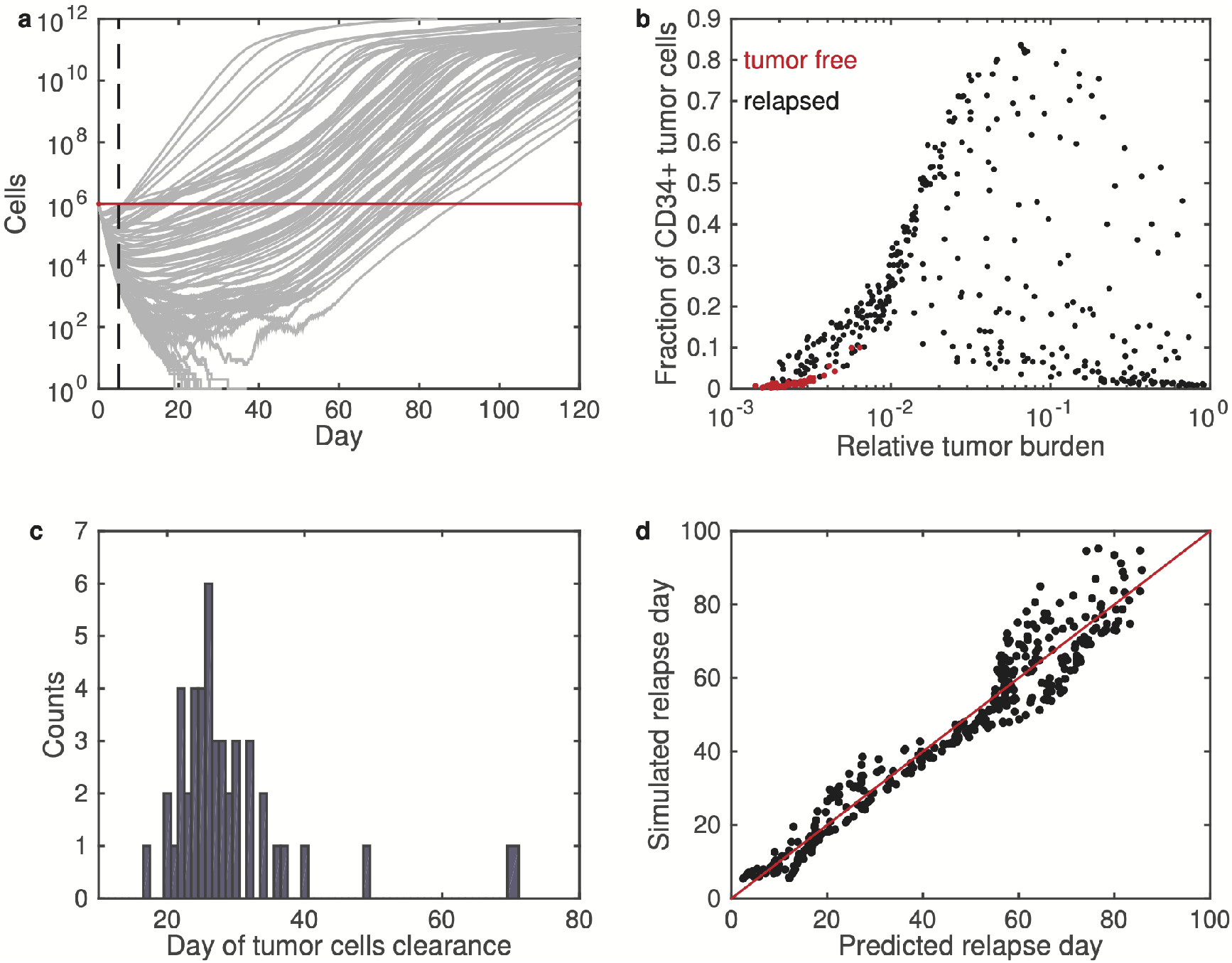
Predictability of the computational model. **a.** Simulated tumor growth (gray lines) after CAR-T cells infusion. The simulation results were obtained by randomly varying the parameters *A*_34_ and *X_0_* over the range 0 < *A*_34_ < 0.8,0 < *X_0_* < 0.6. Black dashed line shows marks day 5 after CAR-T cells infusion, red line shows initial tumor cells number before CAR-T treatment. **b.** Fraction of CD34^+^ tumor cells versus relative tumor burden of all simulated cases in a. **c.** Histogram of the days of tumor cells clearance for the cases with tumor free. **d.** Simulated versus predicted relapse days of all cases with tumor relapse.

For those cases with tumor cells reoccur in the later stage, we defined the timing of tumor relapse as the day when the tumor cell number reached again the initial level after the early remission stage (red line in **Figure 6a**). Data analysis shown that the relapse time (day) can be predicted from the relative tumor burden (*T*) and the fraction of CD34^+^ tumor cell (*f*_CD34_) through a nonlinear function

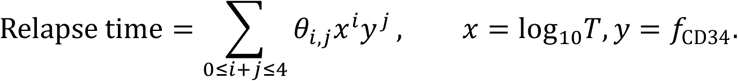

The coefficients *θ_i,j_* were obtained from nonlinear regression based on the data from model simulations. The predicted relapse time shown well agreement with the simulated relapsed time for each case (**Figure 6d)**. Our *in-silico* experiments shown the ability of predicting the outcome of CAR-T treatment through model simulation based on early stage observations of tumor burden and tumor cells analysis.

## Discussions

We developed an individual cell based computational model to study the process of CD19^+^ tumor cells relapse in B-ALL after CD19 targeting CAR-T therapy. The model simulates the collective dynamics of tumor growth in accordance with the kinetic (proliferation, differentiation, or death) rates of individual cells, and highlights the heterogeneity and plasticity at single cell level. In the model, we introduced epigenetic states for each cell, and the kinetic rates of each cell are dependent on the associated epigenetic states. Moreover, the epigenetic states randomly change during cell cycling according to an inheritance probability that was introduced to represent the cell plasticity in response to CAR-T stress. The proposed model enables us to simulate the population dynamics as well as the changes of epigenetic states at individual cells. The model outlines a general computational model framework of simulating collective stem cells regeneration with cell heterogeneous and plasticity. In the model framework, we can overlook detail cellular signaling networks, and focus at how the kinetic rates may depend on the epigenetic states and how the epigenetic states may change during cell cycling.

In our study, to model the process of tumor relapse, we proposed a major hypothesis that CAR-T stresses may induce stem-like cell transition of tumor cells and the immune escape of the stem-like cells. Flow cytometry analysis of tumor cells from relapsed mice shown the existence of CD19^+^CD34^+^ cells that were not presented in the non-treated mice. In the model, we introduced CD34 as the marker gene to represent the stemness of tumor cells. Model simulations nicely reproduce the process of tumor relapse, during which hematopoietic-stem like cells emerge due to CAR-T stress after CAR-T cells infusion. The hematopoietic stem-like tumor cells give rise to downstream cell lineages and may lead to mixed-phenotype acute leukemia, which is crucial for the development of personalized therapy.

Tumor relapse is a common issue in refractory B-ALL therapy; however, wet laboratory experiments are incapable of studying the relapse process. The proposed computational model provides an *in-silico* laboratory to investigate this process in detail and to predict the outcome of CAR-T therapy. For example, if we can measure the tumor burden and estimate the fraction of CD34^+^ tumor cells at the early stage after CAR-T cells infusion, we may turn the model parameters to fit the data and predict long-term responses according to modeling simulations. Moreover, by varying the model conditions, we are able to study other issues of immunotherapy, such as the effects of memory CAR-T cell efficiency(49) and CD19 loss on tumor relapse(11), as well as the dynamical systems perspective on CAR-T cell dosing(50). To the best of our knowledge, this is the first computational model for CAR-T therapy that combines gene expression variations at the single cell level with population dynamics, and that is capable of investigating cell plasticity in response to drug stress. Combinations between *in vivo* experiments and *in silico* simulation can lead to the quantitative design of an optimal strategy and the prediction of therapy effects, which are important for personalized medicine in individual patients.

## AUTHOR CONTRIBUTIONS

**Conception and design:** X. Zhong, J. Lei

**Development of methodology:** X. Zhong, C.Zhang, R. Chang, Y. Bai, J. Lei, X. Jiao

**Acquisition of data:** C. Zhang, Y. Bai, M. Li, H. Shi

**Analysis and interpretation of data:** Y.Bai, C. Zhang, X. Jiao

**Writing, review, and/or revision of the manuscript:** X. Zhong, J. Lei

**Administrative, technical, or material support**: X. Zhong, C. Zhang, Y. Bai, H. Shi

**Study supervision:** X. Zhong, J. Lei

## Acknowledgements

The authors acknowledge Dr. Gattinoni Luca and Michael Sadelain for reading the manuscript and helpful suggestions. This work was support by the Foundation of Beijing Municipal Science and Technology (Z161100000216136) to X. Zhong, and the National Natural Science Foundation of China (NSFC 91730301 and 11831015) to J. Lei.

## Supplemental Figures

**Figure S1.**
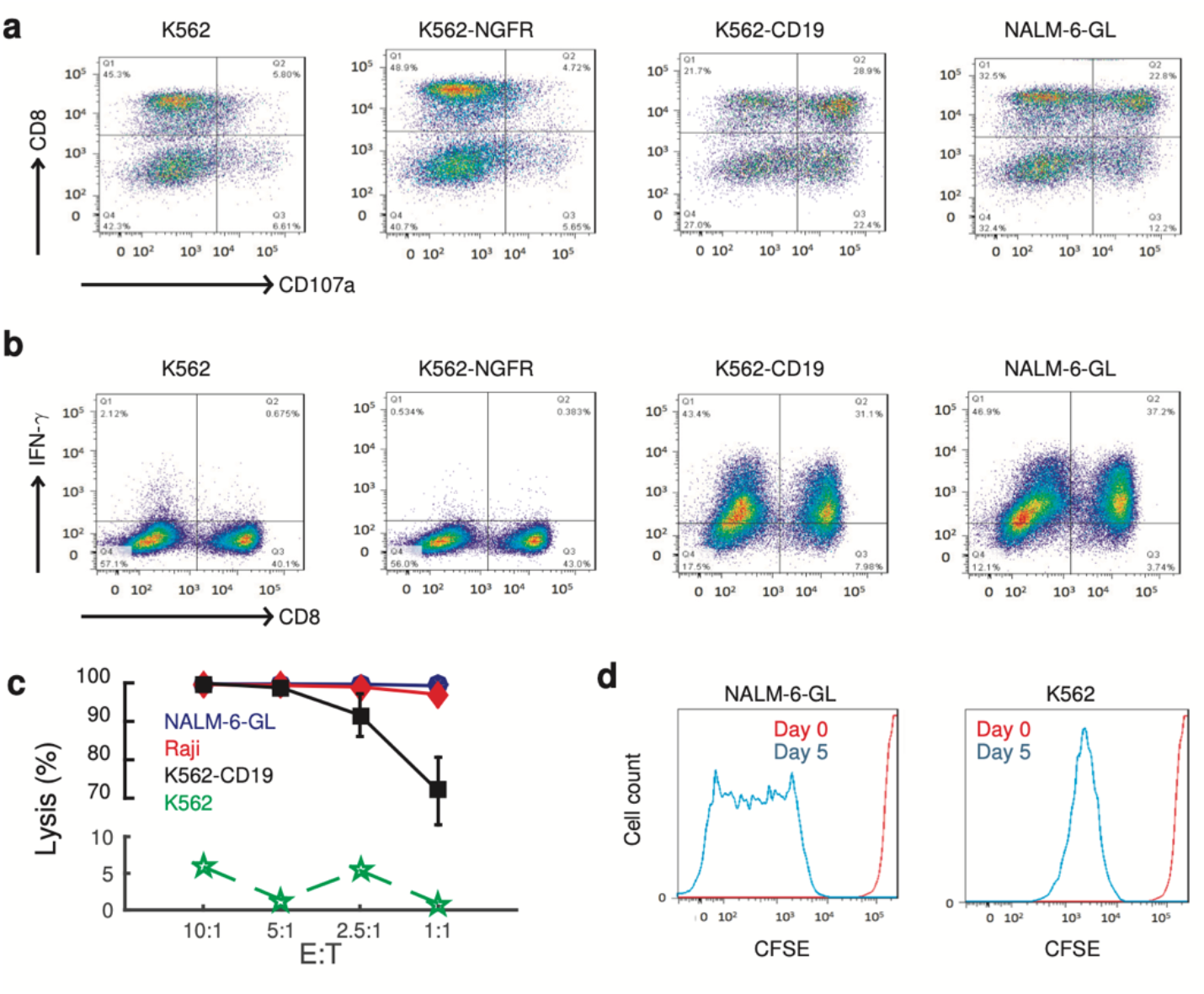
Functional assessment of CD19-28z CAR-T cells. **a.** Flow cytometry plots showing CD107a expression in the CD19-28z CAR-T cells co-cultured with K562, K562-NGFR, K562-CD19, or NALM-6-GL pre-B-ALL cells. **b.** Flow cytometry plots showing IFN-γ expression in CD19-28z CAR-T cells co-cultured with K562, K562-NGFR, K562-CD19, or NALM-6-GL cells. **c.** The lysis percentages of various types of cells co-cultured for 24h with CD19-28z CAR-T cells. **d.** CFSE plots showing the proliferation of the CD19-28z T cells co-cultured with NALM-6-GL or K562 cells at days 0 and 5.

**Figure S2.**
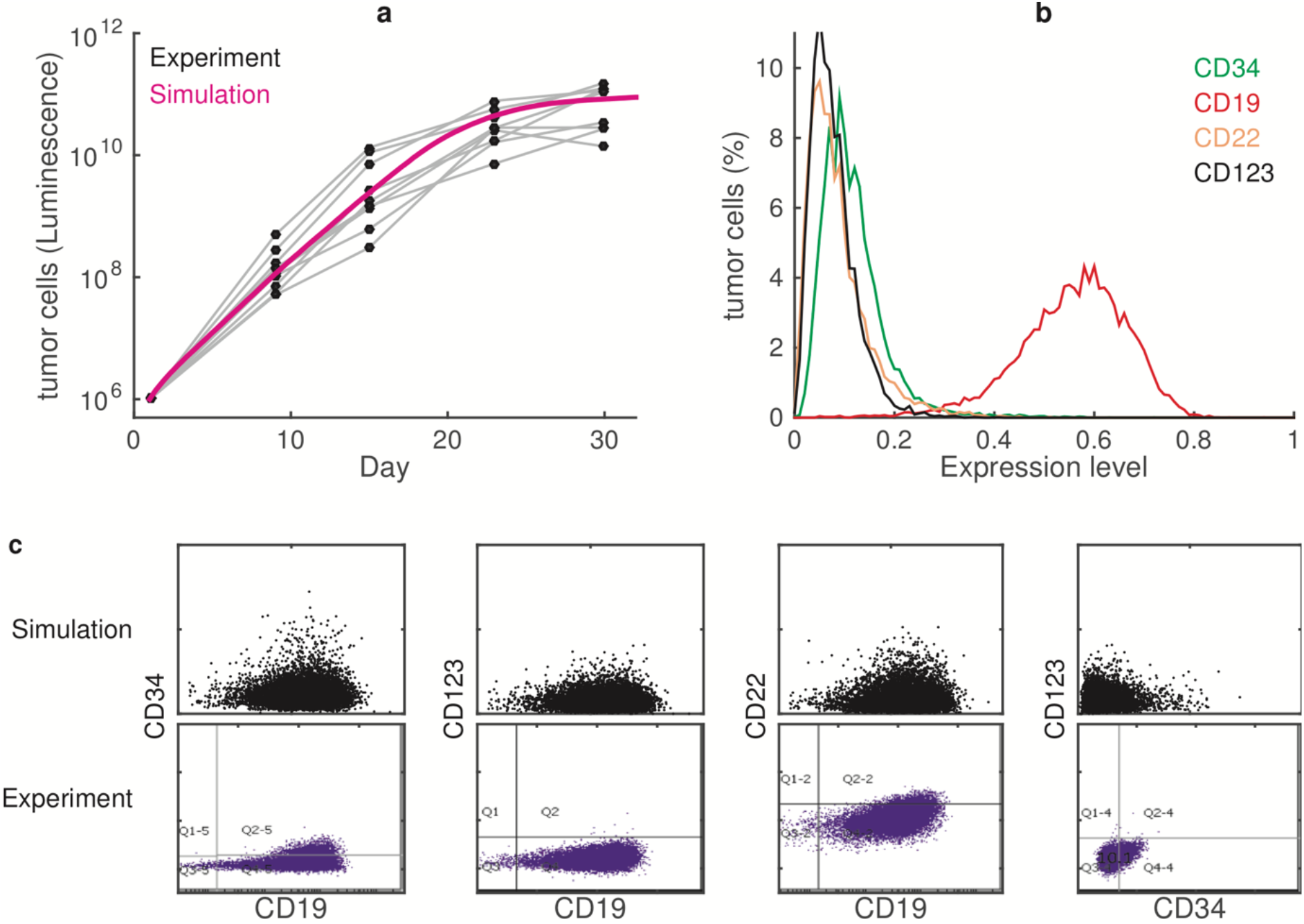
Simulation results with control NGFR-28z T cells. **a.** Evolution of tumor cell numbers (rescaled to compare with the BLI data). The solid lines from the model simulation, and the black dots are from BLI of mice injected with NALM-6-GL cells. **b**. Distribution of the simulated marker expression levels at day 30. **c**. Scatter plots of the marker gene expression levels at day 30, from the simulation (upper panel) and from the flow cytometry analysis (bottom panel).

**Figure S3.**
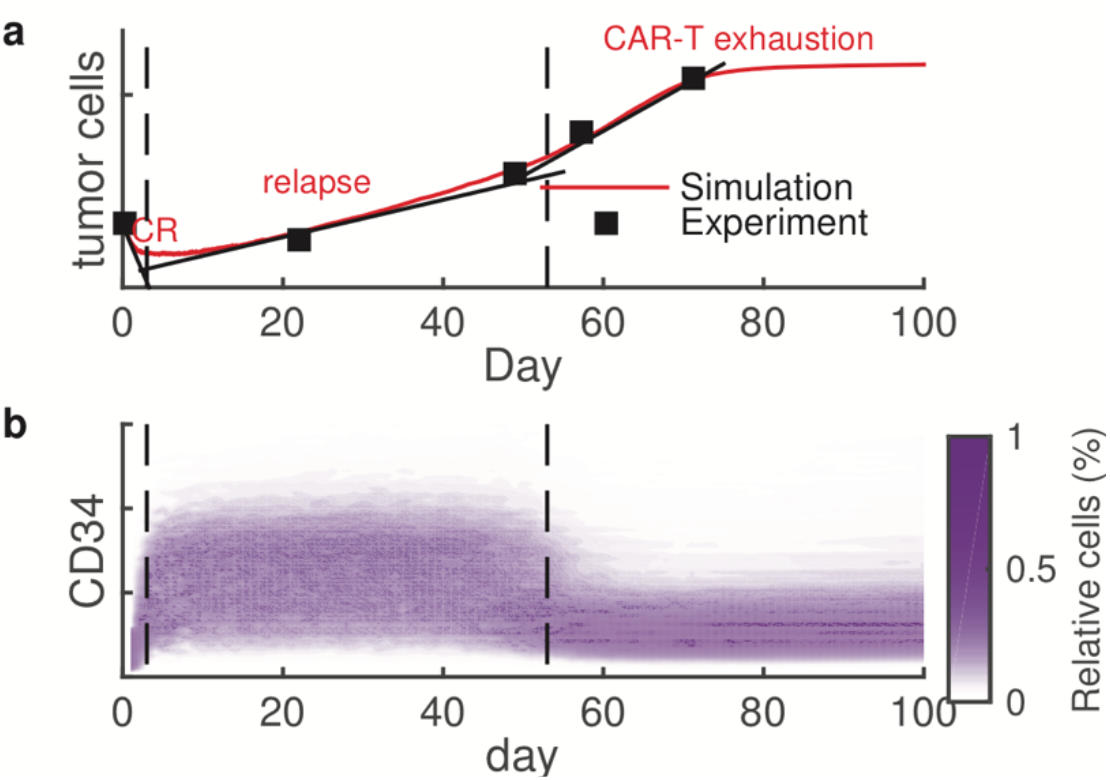
Dynamics of tumor relapse after the CAR-19 T cell treatment. **a.** Three stages of tumor relapse. **b.** Evolution of the density of the relative CD34 expression in tumor cells after the CAR-T cell infusion.

## Supplementary Text

### S1 S1 Experimental Protocols

#### Cell culture and antibodies

NALM-6 pre-B-ALL cells were obtained from ATCC. NALM-6-GFP-Luc cells (NALM-6-GL cells stably transfected with the gene for green fluorescent protein (GFP) and luciferase) were cultured in RPMI 1640 medium supplemented with 10% heat-inactivated foetal bovine serum (FBS) (Gibco), 50 U/ml penicillinstreptomycin (Gibco), 2 mM GlutaMAX (Lonza), and 1 mM sodium pyruvate (Lonza). The PG-13 and Phoenix ECO retroviral producer cell lines were cultured in DMEM supplemented with 10% FBS (Gibco), 50 U/ml penicillinstreptomycin (Gibco), 2 mM GlutaMAX (Lonza), and 1 mM sodium pyruvate (Lonza). T cells were always cultured in T cell medium, which consisted of X-VIVO-15 medium supplemented with 5% AB serum (Sigma-Aldrich), 100 U/ml IL-2, and 50 U/ml penicillinstreptomycin (Gibco). All cells were routinely tested for mycoplasma and found to be negative. The following antibodies were used: anti-CD3-PE (555340, BD Biosciences); anti-CD4-BV421 (562424, BD Biosciences); anti-CD8-Alex Fluor 750 (MHCCD0822, Invitrogen); anti-CD19-PE-Cy7 (SJ25C1, BD Biosciences); anti-CD123-APC (560087, BD Biosciences); anti-CD20-V450 (561164, BD Biosciences); anti-CD22-PerCP-Cy5.5 (561441, BD Biosciences); and anti-CD34-Alexa Fluor 700 (561441, BD Bioscience).

#### Immunohistochemistry analysis

Sections of the obtained bone marrow were analyzed by immunohistochemistry (anti-human CD19 staining) after the NALM-6-GL tumor was explanted. Images were acquired at room temperature using a Nikon Ci Eclipse microscope system (Nikon) with a Nikon Plan Apo VC 20/0.75 objective lens, Nikon DigiSight Digital Camera Head and Nikon NSI-Elements SF 46000 software. Representative regions at 20 magnification were shown.

#### Retroviral transduction

All peripheral blood samples were obtained after volunteers had provided written informed consent under an institutional review board-approved protocol. Retroviral supernatant was generated from the CD19-28z PG-13 Producer Cell Clone as previously described[1] and was collected at 24 and 48 hours. Peripheral blood mononuclear cells from healthy donors were isolated by Ficoll gradient centrifugation (GE Healthcare) and activated with anti-CD3/CD28 T cell Activator Dynabeads (Invitrogen) immediately after purification at a 1:1 bead-to-cell ratio. After 48 hours of bead activation, the T cells were transduced with the retroviral supernatants by centrifugation on Retronectin (Takara)-coated plates to obtain CD19-28z CAR T cells. The transduction efficiency was verified 7 days later by flow cytometry (FITC-conjugated goat anti-mouse IgG(H+L)) (F2653, Sigma-Aldrich). The CAR T cells were injected into mice 12 days after the first T cell activation.

#### Mice

Mice were treated under a protocol approved by the animal ethics committee of Beijing Shijitan Hospital, Capital Medical University. All relevant animal use guidelines and ethical regulations were followed. Female NOD-SCID mice were purchased from the Charles River Laboratories (Beijing, China) and maintained under pathogen-free conditions. After adaptive feeding for 1 week, the mice were intravenously injected on day 1 with 1 × 10^6^ NALM-6-GFP-Luc (NALM-6-GL) cells in 0.2 mL of RPMI 1640 medium, and on days 2, 3, 4 and 12, the mice were injected with 5 × 10^6^ CD19-28z, CD123-BBz or control NGFR-28z T cells daily. In all experiments, the mice that developed hind limb paralysis or decreased responsiveness to stimuli were sacrificed for flow cytometry analysis. Bioluminescence imaging utilized a Xenogen IVIS Imaging System (Xenogen) with Living Image software (Xenogen) for the acquisition of imaging datasets. Both dorsal and ventral views were obtained for all animals. Tumor burden was assessed as previously described[2].

### S2 Model framework

The computational model was developed to simulate the dynamical response to CD19 chimeric antigen receptor (CAR) T cell therapy for acute B lymphoblastic leukemia (B-ALL). In the model, we mimic the process of CD19 CAR-T treatment to mouse injected with B-ALL tumor cells (NALM-6-GL), and consider the tumor cells population dynamics as well as the cell plasticity in response to CAR-T treatment.

In the model, we mainly consider the dynamics of tumor cells produced by the injected NALM-6-GL that are essential for the study, and neglect the normal hematopoietic cells. Base on the cell lineage of hematopoietic and progenitor stem cells (HPSCs) [3], all tumor cells in the pool under considered are classified into three subpopulations, which are marked by their CD34 expression levels: the hematopoietic stem cell like (HSC-like) cells with high CD34 expression, the progenitors (either myeloid or lymphoid) with intermediate CD34 expression, and terminal differentiated (TD) cells with extreme low CD34 expression. The three population cells different from each other by their dynamical properties: HSC-like cells have low proliferation rate, and can escape CAR-T treatment; progenitors cells have high proliferation rate, can differentiate to TD cells, and CD19 positive lymphoid progenitor cells can be killed by CD19 CAR-T; the TD cells are not renewable and can actively be destroyed from the body. We further assume that the expression levels of marker genes change during cell division, which result in the plasticity of tumor cells (to be detailed below). In comparing with experimental data, we mainly considered renewable cells (HPSC-like cells).

The developed model is single-cell based, in which a pool of renewable hematopoietic and progenitor cells are considered, each cells is described individually through its own epigenetic state; the epigenetic state of each cells is dynamically changed during cell regeneration. The model only consider tumor cells in the body, which are originally injected into the mice through blood injections. Normal cells are not considered explicitly. Moreover, only self-renewal cells are considered in the model, the terminal differentiated cells that loss the ability of self-renewal are not included explicitly. The population dynamics of tumor cells including proliferation, apoptosis, and removal through CAR-T cells are modeled with single cell stochastic behaviors.

In the model, each cell is represented by the expression of marker genes CD19, CD34, CD22, CD123. Here, CD34 is a marker of HSCs, and can be used to present the ability of self-renew (proliferation rate)[4, 5, 6]. CD19 is a marker of CAR-T target, and represent the ability of being recognized and killed by CAR-T cells (CD19-28z). CD22 expression can be considered as mostly constant in precursor B-cells, and can be promoted in CD19+ Pro-B cells under stress with CD19 CAR-T[7]. Moreover, CD22 can block the effect of CAR-T to CD19, and induce the homing of recirculating B cells[8]. CD123 (IL3RA) encodes an interleukin 3 specific subunit of a heterodimeric cytokine receptor, which is a marker of myeloid cell line. CD123 expression is often seen in B-ALL patients [9, 10, 11]. **Figure S4** shows the distribution of expression levels of the four marker genes based on single-cell RNA-sequencing for HPSC.

### S3 Major assumptions and formulations

Here, we shown major assumptions and formulations of the model. Sketch of the main assumptions are shown at **Figure 1a-b**, and are listed below:

1. **Epigenetic state of tumor cells** The epigenetic state of each cell is represented by the expressions of CD34 ([CD34]), CD19 ([CD19]), CD22 ([CD22]), and CD123 (CD123]). All expression levels are normalized with respect to their maximum level, respectively, so that the corresponding values vary continuously over the interval from 0 to 1. The continuous distribution of marker gene expressions are supported by single-cell RNA sequencing of HPSC [12, 13]. **Figure S4** shows the distribution of these markers genes based on single-cell RNA-sequencing of 1414 hematopoietic stem and progenitor cells from health persons (1035 cells from a 25-year-old male, and 379 cells from a 29-year-old female) [12]. Here, we note that the stem cell marker CD34 show bimodal distribution for health persons. There are lots of dropouts for genes CD19, CD22, and CD123, which indicate that they are usually low expression in normal HPSC.

**Figure S4:**
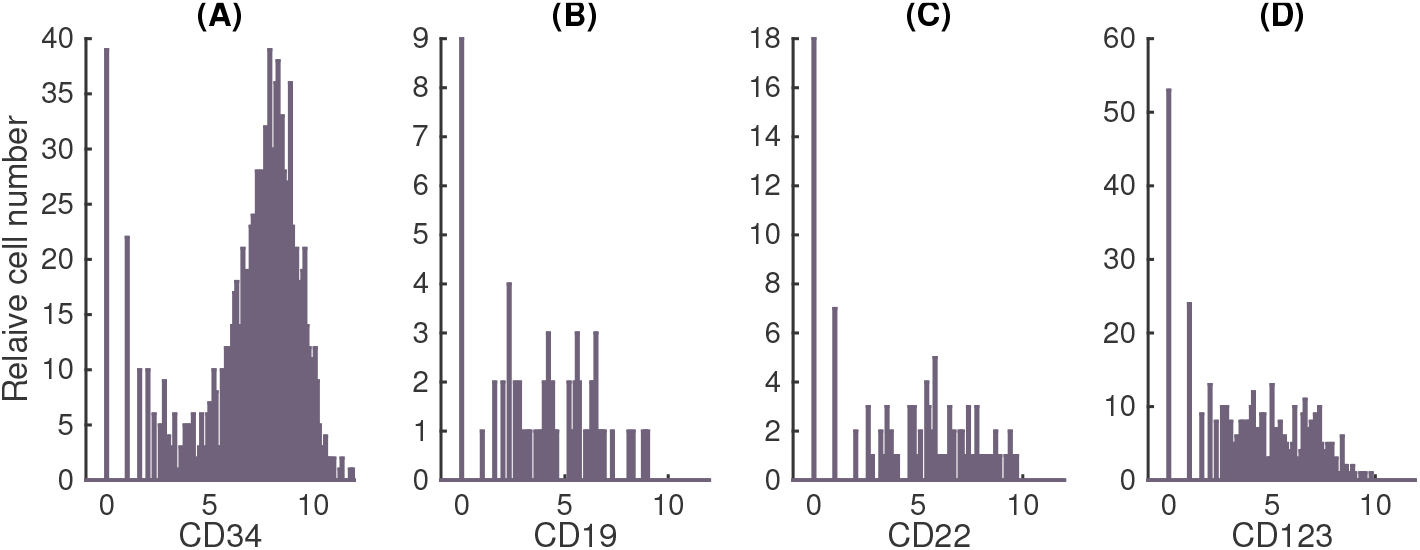
Distribution of the four marker genes CD34, CD19, CD22, and CD123 in hematopoietic and progenitor cells obtained from single-cell RNA-sequencing (GSE75478) [12]. Here we note that in single-cell RNA-seq, events of ‘dropout’ often happen to those genes with low transcription level. Hence, there are a lot of dropout events in low expression genes. In this data set, 217, 1345, 1310, 957 cells, within total 1414 cells, are dropout for genes CD34, CD19, CD22, CD123, respectively.
2. **CAR-T activity** In experiments, CAR-T cells (CD19 18Z) were injected into mice at days 2-4 and 12, respectively, with 5 × 10^6^ cells daily. CFSE tracking show that CAR-T cells are capable of proliferation after being active with NALM-6-GL cells (**Figure S1d**). Immunocytochemistry shown that there are T cells remained at various tissues at weeks 2-5 of the experiments, which show the persistence of T cells at least for 5 weeks. In the model, we do not explicitly model the dynamics of CAR-T activity after injecting into the mice; however we introduce a predefine function *R*(*t*) for the CAR-T activity. We take the function so that *R*(*t*) increase at the early stage, and then exhaust at 7 weeks. Explicitly, we have

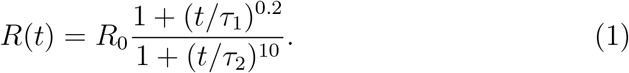 Here, *R*(*t*) first increases from *R* = *R*_0_ (here we set *R*_0_ = 1 by normalizing the CAR-T activity with the initial level) at *t* = 0, and then decreases to 0 when the time t is long enough. The parameters *τ*_1_ and *τ*_2_ (day) are time constants of CAR-T activation and exhaustion, respectively. **Figure S5A** shows the function *R*(*t*) with typical parameters used in model simulation.

**Figure S5:**
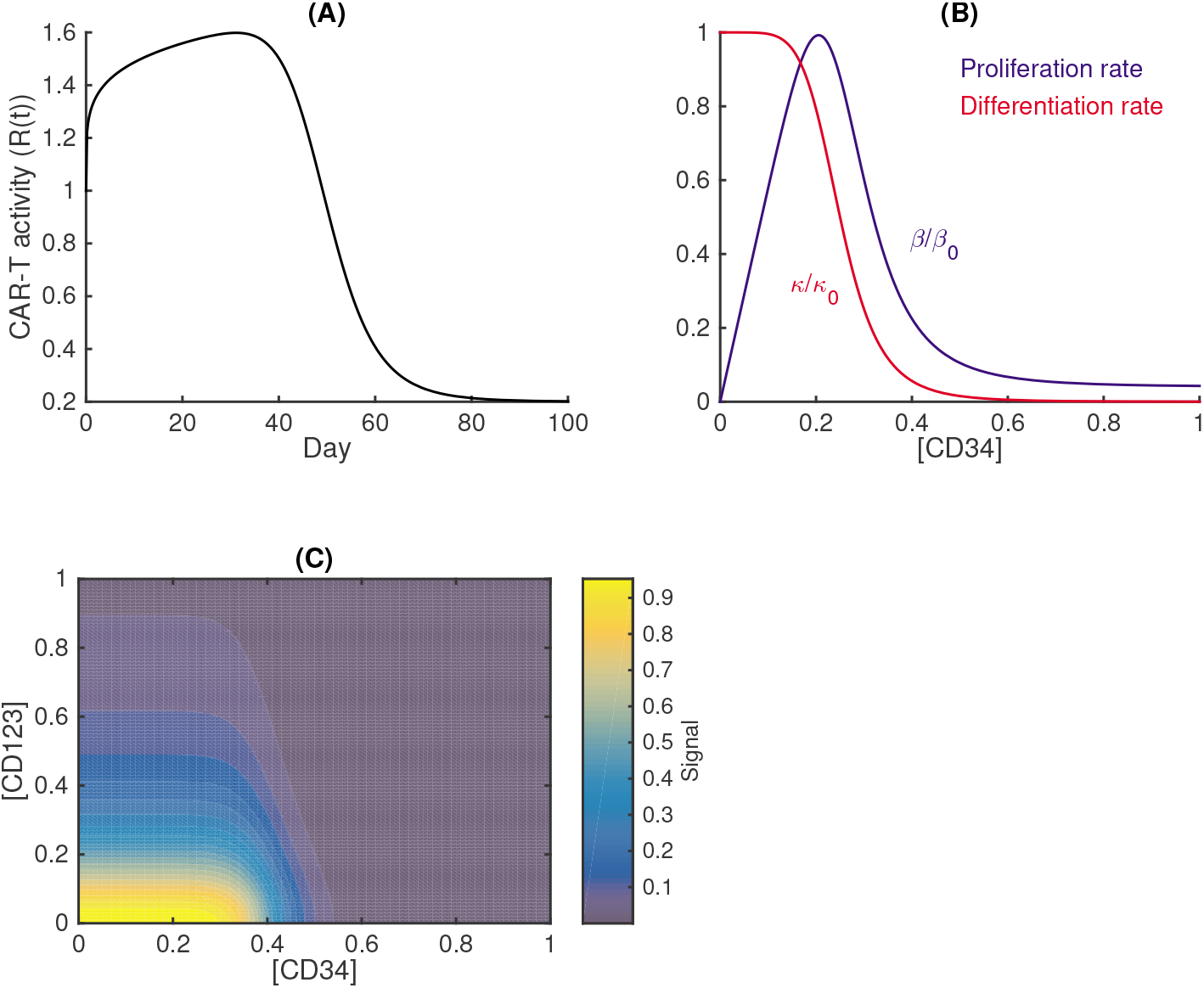
Plots of functions used in the model. (A) The function *R*(*t*) for CART activity (refer Eq. (1)). Here *R*_0_ = 1.0, *τ*_1_ = 230 day, *τ*_2_ = 50 day. (B) The proliferation rate function 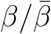 (blue, Eq. (2)), and the terminal differentiation rate function *κ/κ*_0_ (red, refer Eq. (4)). (C) Dependence of CAR-T signal with CD34 and CD123 expression levels (refer Eq. (7)).
3. **Proliferation rate** The proliferation rate *β* (day^-1^) of each cell is dependent on the transcription level of CD34 that represents the stemness of the cell. We formulate the dependence as Hill type function as

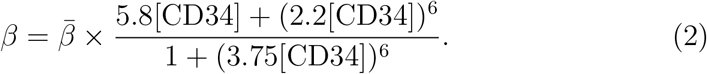 Here 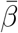 is the maximum proliferation rate with high CD34, which depends on the total cell number due to saturation

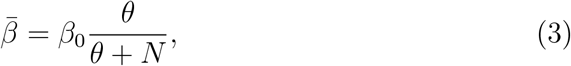

where *N* is the number of tumor cells. **Figure S5B** shows the relative rate *β/β*_0_ (blue curve).
4. **Differentiation rate** All mature precursor cells can under go terminal differentiation with a rate *κ* that depends on the stemness (through CD34) as

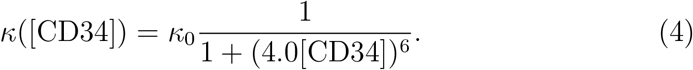
5. **Death rate** Each cell under cell death (either spontaneous apoptosis or due to CAR-T treatment) with a rate *μ* (day^-1^). Mathematically, we write the death rate as

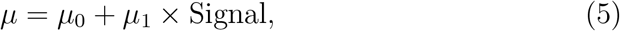

where *μ*_0_ represents the basal apoptosis rate that is independent to CAR-T, and *μ*_1_ is the maximum death rate due to CAR-T treatment, and Signal measures CAR-T signal that is detailed below.
6. **CAR-T signal** The CAR-T signal in tumor cells is determined by the interactions between CAR-T cells and tumor cell surface markers. The interaction is complex and depending on detail reactions in the T cell receptor signaling pathways. Here, we omit the detail reactions, however, focus at the overall dependence of CAR-T signaling with cell types. The signal strength is dependent on the interaction between CD19 and CART, and CD22 expression can repress the interaction, which is formulated as

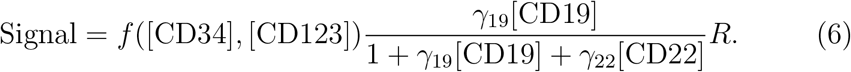 Here *a*_19_ and *a*_22_ are weight coefficients that CD19 and CD22 are recognized by CAR-T. The function *f* represents the signaling factor for the effects of CD34 and CD123 to the CAR-T signal. HSC-like cell (high CD34 expression) is protected from the response to CART, and the role of CD19 CAR-T is limited in myeloid-like cells that expression CD123. Hence, the CAR-T signal is repressed with CD34 and CD123 expression. Hence, we define the signaling factor *f* as

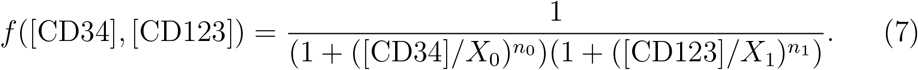 **Figure S5C** shows the function *f* ([CD34], [CD123]).
7. **Epigenetic state transition** Finally, we consider the transition of epigenetic states of each single cell during cell cycling. At each cell cycle, a cell divides into two daughter cells, expressions of [CD34], [CD19], [CD22], and [CD123] in the daughter cells change randomly from the mother cell following a rule of random inheritance. The random inheritance of epigenetic state can be originated from the rearrangement of epigenetic states, *i.e.,* histone modification or DNA methylations [14, 15, 16]. In our previous study of gene expression with epigenetic modifications [17, 18], we have seen that the distribution of modified nucleosomes of daughter cells depends on that of mother cells through binomial distribution. Analogically, we introduce Beta-distribution, a continuous version of the binomial distribution, to describe the cross cell division epigenetic state transition. Moreover, we assume that CAR-T signals can interfere the transitions of epigenetic state over cell cycling. Detail assumption and formulations the markers are given below.

**CD34** Let the transcription level of CD34 at the *k*^th^ cycle as [CD34] = *u_k_* (0 ≤ *u_k_* ≤ 1). After a cell division, the state of two daughter cells 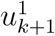 and 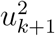 are given below: 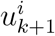 are Beta-distribution random numbers with probability density

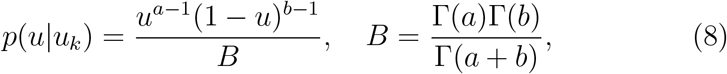

the parameters *a* and *b* are dependent on *u_k_*. For the simplicity, let 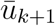 the perspective average level of *u*_*k*+1_ at the (*k* + 1)’th cycle given the expression *u_k_* at the *k*’th cycle, and assume that the mean and variance of *u*_*k*+1_ are

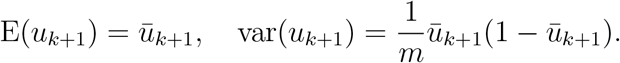

Here *m* is a parameter for the variance (here we take *m* = 60). Then

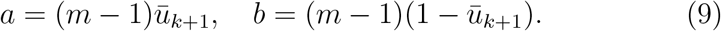

Hence, to obtain the dependence between *u*_*k*+1_ and *u_k_*, we only need to write down how 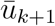 depends on *u_k_*. For CD34, we take

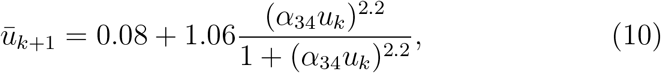

the coefficient *α*_34_ depends on the CD19 expression level and the CAR-T signal

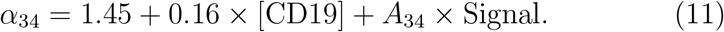

Here, the function (10) is chosen to reproduce the stationary CD34 distribution of injected NALM-6-GL under control NGFR 28Z T cells (**Figure S6**). We can adjust the parameter α_34_ for different stationary distribution of CD34 expression level. The dependence (11) is introduced for the promotion of CD34 expression by CD19 and the CAR-T signaling, with varying coefficient *A*_34_ for the induction of CD34 expression through CAR-T signals.

**Figure S6:**
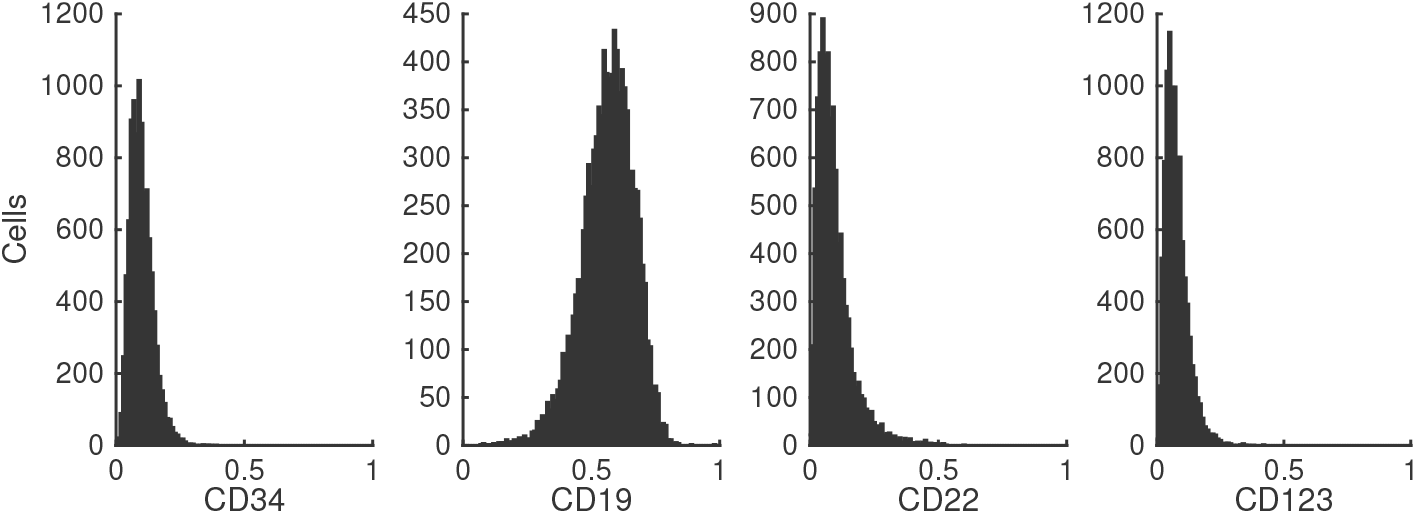
Distributions of simulated marker genes expression of NALM-6-GL cells under control NGFR-28z T cells.

**CD19** The transcription level of CD19 at the *k*^th^ cycle is [CD19] = *v_k_* (0 ≤ *v_k_* ≤ 1). After cell division, the two daughter cells are 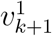 and 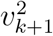, respectively; and 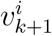 are Beta-distribution random numbers with the probability density function *f* (*v|v_k_*; *a,b*) given by (8), and the parameters *a* and *b* are given by (9) through the mean and variance of *v*_*k*+1_. The function for how the average value of 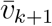 depends on *v_k_* is selected in according to the distribution of CD19 at NALM-6-GL cells, so that

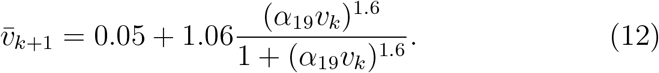

Here, the parameter *α*_19_ can be adjusted for the distribution of different types of progenitor B cells. **Figure S6** shows the simulated stationary distribution of [CD19] in NALM-6-GL cells.

**CD22** The transcription level of CD22 at the *k*^th^ cycle is [CD22] = *w_k_* (0 ≤ *w_k_* ≤ 1). After cell division, the two daughter cells are 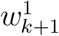 and 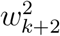, respectively; 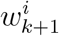 are Beta-distribution random numbers with the probability density function *f* (*w|w_k_*; *a,b*) given by (8), and the parameters *a* and *b* are given by (9) through the mean and variance of *w*_*k*+1_. The average value 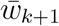 depends on *w_k_* in a way similar to (12). We assume that CD19 promote CD22 expression, and CAR-T signal can also increase the expression of CD22. Hence, we have

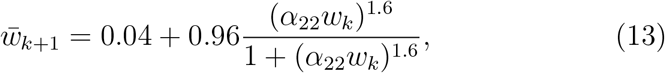

where the coefficient *α*_22_ depends on CD19 expression and the CAR-T signal so that

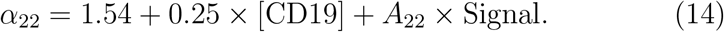

**Figure S6** shows the distribution of CD22 expression in the absence of CAR-T signal.

**CD123** The transcription level of CD123 at the *k*^th^ cycle is [CD123] = *z_k_* (0 ≤ *z_k_* ≤ 1). After cell division, the two daughter cells are 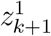 and 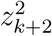, respectively; 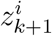 are Beta-distribution random numbers with the probability density function *f* (*z|z_k_*; *a,b*) given by (8), and the parameters *a* and *b* are given by (9) through the mean and variance of *z_k+1_*. The average value 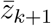 depends on *z_k_* in a way similar to (13), which is given by

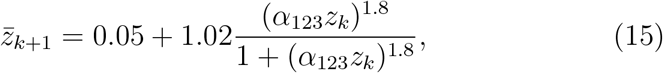

where the parameter *α*_123_ depends on CD34 and the CAR-T signal

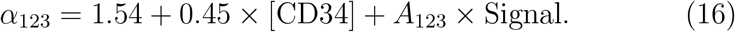

Here we assume that CD34 promote the expression of CD123, which mimic the differentiation from HSC-like to myeloid-like cells. Moreover, we assume that CAR-T signal can also promote the expression of CD123. **Figure S6** shows the distribution of CD123 expression in the absence of CAR-T signal.

### S4 Parameter values

To estimate parameters used in the simulation, we first find the maximum proliferation rate *β*_0_ and basal death rate *μ*_0_ of tumor cells by *in vitro* cell cultures. Next, we adjust the differentiation rate *κ*_0_, and proliferation saturation coefficient *θ*, and other related parameters by comparing simulation results with experimental data under control condition. Finally, we find the parameter values related to CAR-T signaling and the responses by comparing numerical simulation with experimental data.

First, to estimate the proliferation and death rate of tumor cells, we culture the NALM-6-GL cell lines and fit the experimental data with the population dynamics model.

Table S1 shows the cell numbers from day 0 to 3 after cell culturing. The cell number dynamics are modeled with a differential equation

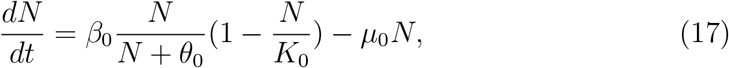

where *N*(*t*) is the cell number at time *t, β*_0_ is the maximum proliferation rate, *θ*_0_ is a parameter associated with growth inhibition, *K*_0_ is a parameter for saturation due to nutrition limitation, and *μ*_0_ is the death rate. Fitting model simulation with experimental data at Table S1, the parameters for the three cell lines are list at Table S2. Comparison between experiment data and simulation is shown at Fig. S7(A). Based on this estimation, we take the maximum proliferation rate of tumors cells as *β*_0_ = 0.12h^-1^ (slightly less than that in cell culturing), and the basal death rate as *μ*_0_ = 8.3 × 10^-5^h^-1^.

**Figure S7:**
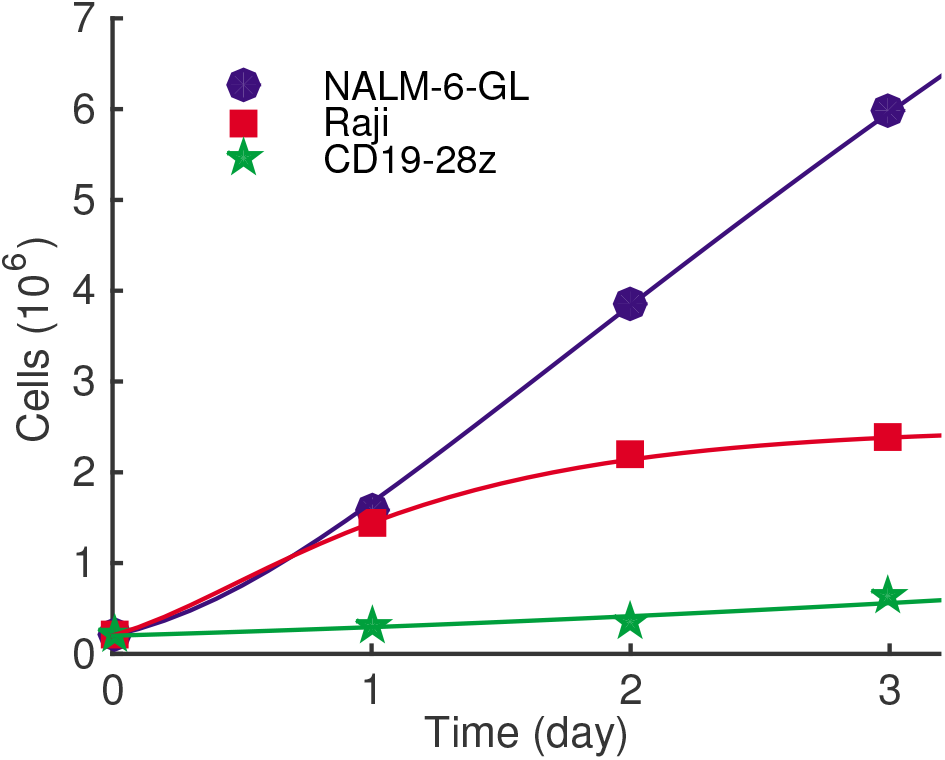
Cell growth dynamics. Markers are experimental data, solid lines are simulation results based on the equation g(17). In simulations, initial condition is *N*(0) = 0.2, and parameters are listed in Table S2.

Next, we fit the experimental data to identify the differentiation rate *κ*_0_ = 0.058h^-1^ (**Figure S2**). **Figure S2** shows simulation results with control NGFR-28z. Both cell number and the distribution fo marker gene expressions are in good agreement with experimental data.

Parameter values used in model simulation are listed at Table S3 and S4.

**Table S1:** Cell numbers of culturing NALM-6-GL, Raji, and CD19-28z (T cells). Here, cell numbers are in 10^6^ cells per hole.

|  | NALM-6-GL | Raji | CD19-28z |
| --- | --- | --- | --- |
| Day 0 | 0.2 | 0.2 | 0.2 |
| Day 1 | 1.57 | 1.45 | 0.30 |
| Day 2 | 3.84 | 2.18 | 0.37 |
| Day 3 | 5.99 | 2.37 | 0.63 |

**Table S2:** Cell growth parameter values for the three cell lines.

| Cell line | $\beta_0$ ($\text{h}^{-1}$) | $\theta_0$ ($10^6$) | $K_0$ ($10^6$) | $\mu_0$ ($\text{h}^{-1}$) |
| --- | --- | --- | --- | --- |
| NALM-6-GL | 0.1375 | 0.8 | 20 | $8.3 \times 10^{-5}$ |
| Raji | 0.15 | 0.6 | 2.5 | $8.3 \times 10^{-5}$ |
| CD19-28z | 0.01875 | 0.85 | 5.0 | $2.08 \times 10^{-2}$ |

**Table S3:** Default parameter values.

| Parameter | Description | Value | Unit |
| --- | --- | --- | --- |
| $\beta_0$ | Maximum proliferation rate | 0.12 | $\text{h}^{-1}$ |
| $\mu_0$ | Basal death rate | $8.3 \times 10^{-5}$ | $\text{h}^{-1}$ |
| $\mu_1$ | CD19 CAR-T induced death rate | 0.12 | $\text{h}^{-1}$ |
| $\kappa_0$ | Differentiation rate | 0.05 | $\text{h}^{-1}$ |
| $\theta$ | Proliferation saturation coefficient | $5.0 \times 10^5$ | $10^6$ cells |
| $X_0$ | Inhibition coefficient of CAR-T by CD34 | $[0, 0.6]^{(a)}$ | — |
| $n_0$ | Hill coefficient | 10 | — |
| $X_1$ | Inhibition coefficient of CAR-T by CD123 | 0.2 | — |
| $n_1$ | Hill coefficient | 2 | — |
| $\gamma_{19}$ | coefficient of CD19 in CAR-T signal | 3.0 | — |
| $\gamma_{22}$ | coefficient of CD22 in CAR-T signal | 1.0 | — |
| $\gamma_{123}$ | coefficient of CD123 in CAR-T signal | 1.5 | — |
| $\alpha_{19}$ | CD19 regulation coefficient | 1.73 | — |
| $A_{22}$ | Effective coefficient of CAR-T signal to CD22 | 0.1 | — |
| $A_{123}$ | Effective coefficient of CAR-T signal to CD123 | 0.20 | — |
| $A_{34}$ | Effective coefficient of CAR-T signal to CD34 | $[0, 0.8]^{(a)}$ | — |

**Table S4:** Parameters used for the mice #1-#5 in **Figure 4a**.

| Mouse | $A_{34}$ | $X_0$ |
| --- | --- | --- |
| #1 | 0.66721 | 0.2814 |
| #2 | 0.54208 | 0.2403 |
| #3 | 0.48216 | 0.2195 |
| #4 | 0.18339 | 0.4275 |
| #5 | 0.12093 | 0.4476 |

### S5 Numerical scheme

In simulations, we start from *N*_0_ = 10^6^ cells, each with initial condition of high CD19 (0.6 < [CD19] < 1.0), low CD34 (0.01 < [CD34] < 0.1), low CD123 (0.01 < [CD123] < 0.1), and low CD22 (0.01 < [CD22] < 0.1). We took the time step Δ*t* = 0.25h to simulate the model dynamics. Each cell in the pool proliferates, dies, or differentiates randomly and independently according the rate function defined by the above formulations; when a cell proliferates, it produces two daughter cells, each has their own marker gene expression levels according to the rules of epigenetic state transition. The sketch of the numerical scheme is given below.

**Initialize** the time *t* = 0, the cell number *Q*, and all cells 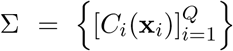. Here **x** represents the vector of epigenetic states ([CD19], [CD34], [CD22], [CD123]).

**for t** from 0 to *T* with step Δ*t* **do**

**for** all cells in Σ **do**

† Calculate the proliferation rate *β*, the apoptosis rate *μ*, and the terminate differentiation rate *κ*.

† Determine the cell fate during the time interval (*t,t* + Δ*t*): The cell is removed (through apoptosis) with a probability *μ*Δ*t*, or undergo terminal differentiation with a probability *κ*Δ*t*, or divides into two cells with a probability *β*Δ*t*.

† If the cell undergo cell division, it is replaced by two daughter cells, and the epigenetic state of each daughter cell is determined according to the inheritance probability function *p*(**x|y**).

**end for**

**Update** the system Σ with the cell number, epigenetic states of all surviving cells, and the ages of the proliferating phase cells, and set *t* = *t* + Δ*t*.

**end for**

The tumor cells number can increase to a high number of 10^12^, which may cause a challenge issue in simulations while we simulate and store each cell. To overcome this issue, we applied a method of *sub-culturing*. We predefined a maximum number of cells to be simulated and stored (*N*_max_ = 10^6^ cells in the current simulation). Let *N_k_* the number of cells under simulation at step *k*, we first open a storage space for 2*N_k_* cells (the maximum number of cells if all *N_k_* cells are divided). After performing cell fate decision for each cell, we have potentially *N*_temp_ (*N*_temp_ ≤ 2*N_k_*) cells. If *N*_temp_ > *N*_max_, we randomly select at most *N*_max_ cells (select each cells with a probability *p* = *N*_max_ < *N*_temp_, and totally no more than *N*_max_ cells) to obtain *N*_next_ cells for the next step simulation. Finally, we store the state of all selected *N*_next_ cells, let *f*_pro,*k*_ = *N*_temp_/*N_k_* for the proliferation rate, and *N*_*k*+1_ = *N*_next_ for the cell number simulated at the next step.

According the approach of sub-culturing simulation, at step *k*, there are *N_k_* ≤ *N*_max_ cells under simulation, and the states of these cells are stored; the real cell number in tumor is given by

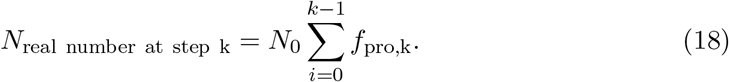

This gives the tumor cells number, which is to be compared with the luminescence (after scale a factor) from experiments.

### S6 Prediction of the day of tumor relapse

To examine the predictability of the computational model, we varied the parameters *A*_34_ and *X*_0_ over a range of 0 < *A*_34_ < 0.8,0 < *X*_0_ < 0.6. For each parameter pair, we simulated the model and obtained the day of tumor relapse as the timing when the tumor cell number reached agin the initial level before CAR-T infusion. Moreover, for each case, we calculated the relative tumor burden (*T*) and the fraction of CD34^+^ cells *f*_CD34_ at day 5 after CAR-T infusion. Here, we only considered the cases with early stage remission, and hence the relative tumor burden *T* < 1.

Based on the simulation data, we find the fitting of relapse time with a nonlinear function

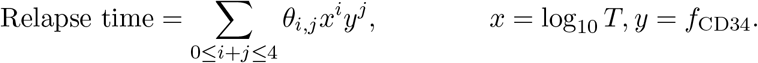

The coefficients *θ_i;j_* we obtained by nonlinear regression through the function `FindFit` in Mathematica. The program and results of fitting are shown at **Figure S8**. The fitting result gives the coefficients

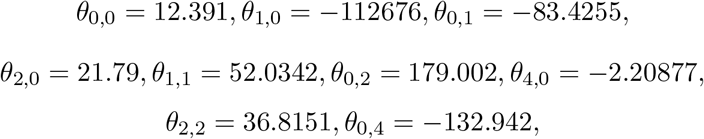

and other coefficients are zero.

**Figure S8:**
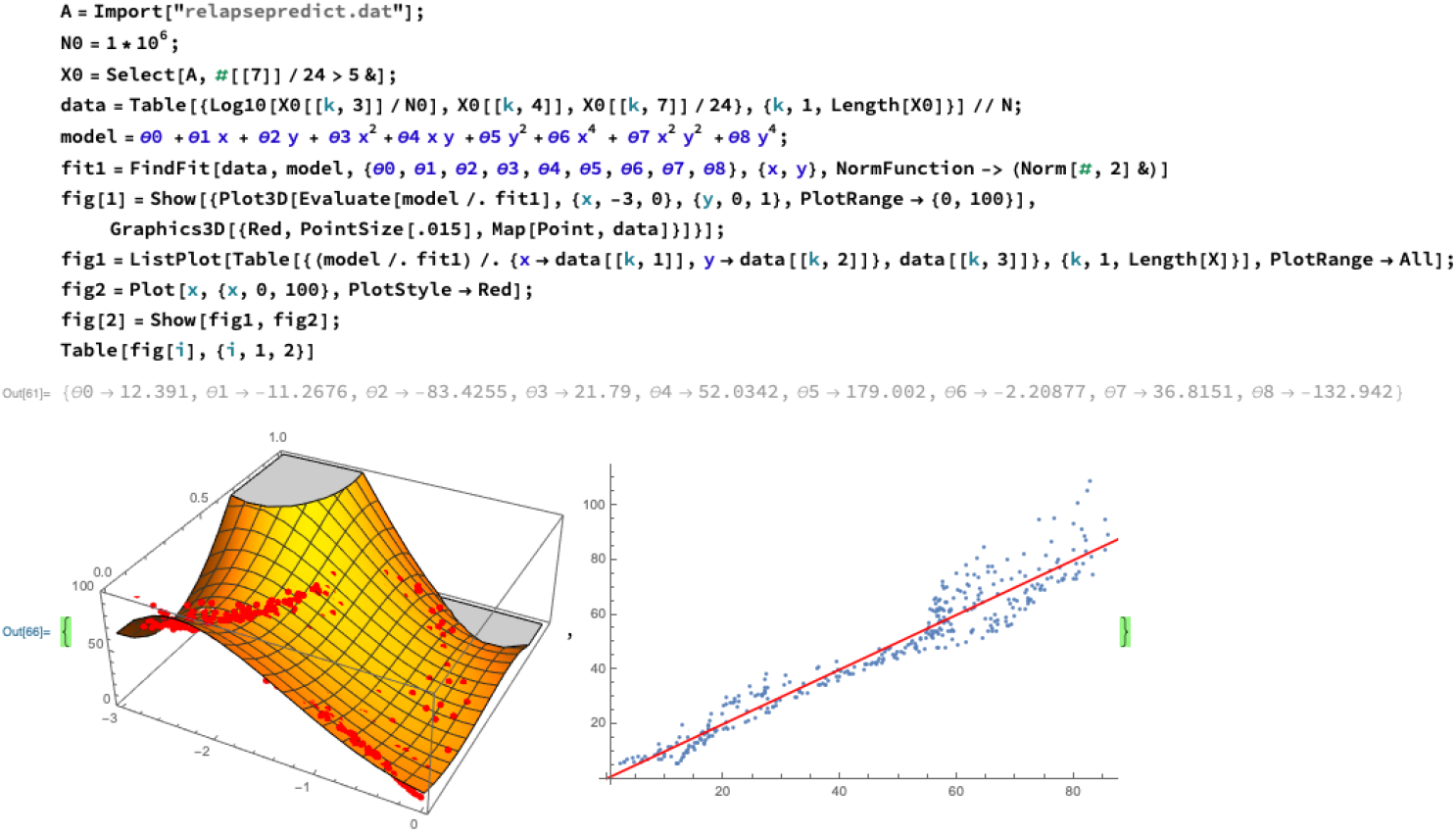
Fitting of the relapse time through Wolfram Mathematica 12.

## REFFERENCES

1. June CH, O’Connor RS, Kawalekar OU, Ghassemi S, Milone MC. CAR T cell immunotherapy for human cancer. Science. American Association for the Advancement of Science; 2018;359:1361–5.

2. Gardner RA, Finney O, Annesley C, Brakke H, Summers C, Leger K, et al. Intent-to-treat leukemia remission by CD19 CAR T cells of defined formulation and dose in children and young adults. Blood. American Society of Hematology; 2017;129:3322–31.

3. Fraietta JA, Nobles CL, Sammons MA, Lundh S, Carty SA, Reich TJ, et al. Disruption of TET2 promotes the therapeutic efficacy of CD19-targeted T cells. Nature. 2018;558:307–12.

4. Labanieh L, Majzner RG, Mackall CL. Programming CAR-T cells to kill cancer. Nat Biomed Eng. Nature Publishing Group; 2018;2:377–91.

5. June CH, Sadelain M. Chimeric Antigen Receptor Therapy. N Engl J Med. 2018;379:64–73.

6. Shah NN, Fry TJ. Mechanisms of resistance to CAR T cell therapy. Nat Rev Clin Oncol. Nature Publishing Group; 2019;3:95ra73–385.

7. Giavridis T, van der Stegen SJC, Eyquem J, Hamieh M, Piersigilli A, Sadelain M. CAR T cell-induced cytokine release syndrome is mediated by macrophages and abated by IL-1 blockade. Nat Med. 2018;24:731–8.

8. Brudno JN, Kochenderfer JN. Chimeric antigen receptor T-cell therapies for lymphoma. Nat Rev Clin Oncol. 2018;15:31–46.

9. Wang J, Hu Y, Huang H. Acute lymphoblastic leukemia relapse after CD19-targeted chimeric antigen receptor T cell therapy. J Leukoc Biol. 2017;102:1347–56.

10. Ruella M, Maus MV. Catch me if you can: Leukemia Escape after CD19-Directed T Cell Immunotherapies. Comput Struct Biotechnol J. 2016;14:357–62.

11. Majzner RG, Mackall CL. Tumor Antigen Escape from CAR T-cell Therapy. Cancer Discov. 2018;8:1219–26.

12. Park JH, Rivière I, Gonen M, Wang X, Sénéchal B, Curran KJ, et al. Long-Term Follow-up of CD19 CAR Therapy in Acute Lymphoblastic Leukemia. N Engl J Med. 2018;378:449–59.

13. Maude SL, Laetsch TW, Buechner J, Rives S, Boyer M, Bittencourt H, et al. Tisagenlecleucel in Children and Young Adults with B-Cell Lymphoblastic Leukemia. N Engl J Med. 2018;378:439–48.

14. Locke FL, Neelapu SS, Bartlett NL, Siddiqi T, Chavez JC, Hosing CM, et al. Phase 1 Results of ZUMA-1: A Multicenter Study of KTE-C19 Anti-CD19 CAR T Cell Therapy in Refractory Aggressive Lymphoma. Mol Ther. 2017;25:285–95.

15. Rossig C, Pule M, Altvater B, Saiagh S, Wright G, Ghorashian S, et al. Vaccination to improve the persistence of CD19CAR gene-modified T cells in relapsed pediatric acute lymphoblastic leukemia. Leukemia. 2017;31:1087–95.

16. Turtle CJ, Hanafi L-A, Berger C, Hudecek M, Pender B, Robinson E, et al. Immunotherapy of non-Hodgkin’s lymphoma with a defined ratio of CD8+ and CD4+ CD19-specific chimeric antigen receptor-modified T cells. Sci Transl Med. 2016;8:355ra116.

17. Kebriaei P, Singh H, Huls MH, Figliola MJ, Bassett R, Olivares S, et al. Phase I trials using Sleeping Beauty to generate CD19-specific CAR T cells. J Clin Invest. 2016;126:3363–76.

18. Bhoj VG, Arhontoulis D, Wertheim G, Capobianchi J, Callahan CA, Ellebrecht CT, et al. Persistence of long-lived plasma cells and humoral immunity in individuals responding to CD19-directed CAR T-cell therapy. Blood. 2016;128:360–70.

19. Turtle CJ, Hanafi L-A, Berger C, Gooley TA, Cherian S, Hudecek M, et al. CD19 CAR-T cells of defined CD4+:CD8+ composition in adult B cell ALL patients. J Clin Invest. 2016;126:2123–38.

20. Fraietta JA, Beckwith KA, Patel PR, Ruella M, Zheng Z, Barrett DM, et al. Ibrutinib enhances chimeric antigen receptor T-cell engraftment and efficacy in leukemia. Blood. American Society of Hematology; 2016;127:1117–27.

21. MD DWL, MD JNK, MD MS-S, MD YKC, RN CD, PhD SAF, et al. T cells expressing CD19 chimeric antigen receptors for acute lymphoblastic leukaemia in children and young adults: a phase 1 dose-escalation trial. Lancet. Elsevier Ltd; 2015;385:517–28.

22. Kochenderfer JN, Dudley ME, Kassim SH, Somerville RPT, Carpenter RO, Stetler-Stevenson M, et al. Chemotherapy-refractory diffuse large B-cell lymphoma and indolent B-cell malignancies can be effectively treated with autologous T cells expressing an anti-CD19 chimeric antigen receptor. Journal of Clinical Oncology. 2015;33:540–9.

23. Kochenderfer JN, Dudley ME, Carpenter RO, Kassim SH, Rose JJ, Telford WG, et al. Donor-derived CD19-targeted T cells cause regression of malignancy persisting after allogeneic hematopoietic stem cell transplantation. Blood. American Society of Hematology; 2013;122:4129–39.

24. Cruz CRY, Micklethwaite KP, Savoldo B, Ramos CA, Lam S, Ku S, et al. Infusion of donor-derived CD19-redirected virus-specific T cells for B-cell malignancies relapsed after allogeneic stem cell transplant: a phase 1 study. Blood. American Society of Hematology; 2013;122:2965–73.

25. Schuster SJ, Svoboda J, Chong EA, Nasta SD, Mato AR, Anak Ö, et al. Chimeric Antigen Receptor T Cells in Refractory B-Cell Lymphomas. N Engl J Med. 2017;377:2545–54.

26. Kalos M, Levine BL, Porter DL, Katz S, Grupp SA, Bagg A, et al. T cells with chimeric antigen receptors have potent antitumor effects and can establish memory in patients with advanced leukemia. Sci Transl Med. American Association for the Advancement of Science; 2011;3:95ra73–3.

27. Fischer J, Paret C, Malki El K, Alt F, Wingerter A, Neu MA, et al. CD19 Isoforms Enabling Resistance to CART-19 Immunotherapy Are Expressed in B-ALL Patients at Initial Diagnosis. J Immunother. Journal of Immunotherapy; 2017;40:187–95.

28. Sotillo E, Barrett DM, Black KL, Bagashev A, Oldridge D, Wu G, et al. Convergence of Acquired Mutations and Alternative Splicing of CD19 Enables Resistance to CART-19 Immunotherapy. Cancer Discov. 2015;5:1282–95.

29. Gardner R, Wu D, Cherian S, Fang M, Hanafi L-A, Finney O, et al. Acquisition of a CD19-negative myeloid phenotype allows immune escape of MLL-rearranged B-ALL from CD19 CAR-T-cell therapy. Blood. American Society of Hematology; 2016;127:2406–10.

30. Perna F, Sadelain M. Myeloid leukemia switch as immune escape from CD19 chimeric antigen receptor (CAR) therapy. Transl Cancer Res. 2016;5:S221–5.

31. Jacoby E, Nguyen SM, Fountaine TJ, Welp K, Gryder B, Qin H, et al. CD19 CAR immune pressure induces B-precursor acute lymphoblastic leukaemia lineage switch exposing inherent leukaemic plasticity. Nat Commun. 2016;7:12320.

32. Oberley MJ, Gaynon PS, Bhojwani D, Pulsipher MA, Gardner RA, Hiemenz MC, et al. Myeloid lineage switch following chimeric antigen receptor T-cell therapy in a patient with TCF3-ZNF384 fusion-positive B-lymphoblastic leukemia. Pediatr Blood Cancer. 2018;:e27265.

33. Torres-Collado AX, Jazirehi AR. Overcoming Resistance of Human Non-Hodgkin’s Lymphoma to CD19-CAR CTL Therapy by Celecoxib and Histone Deacetylase Inhibitors. Cancers (Basel). 2018;10.

34. Graf T. Differentiation plasticity of hematopoietic cells. Blood. 2002;99:3089–101.

35. Laurenti E, Göttgens B. From haematopoietic stem cells to complex differentiation landscapes. Nature. 2018;553:418–26.

36. Takahashi K, Wang F, Morita K, Yan Y, Hu P, Zhao P, et al. Integrative genomic analysis of adult mixed phenotype acute leukemia delineates lineage associated molecular subtypes. Nat Commun. Nature Publishing Group; 2018;9:2670.

37. Gerr H, Zimmermann M, Schrappe M, Dworzak M, Ludwig W-D, Bradtke J, et al. Acute leukaemias of ambiguous lineage in children: characterization, prognosis and therapy recommendations. Br J Haematol. 2010;149:84–92.

38. Matutes E, Pickl WF, Veer MV, Morilla R, Swansbury J, Strobl H, et al. Mixed-phenotype acute leukemia: clinical and laboratory features and outcome in 100 patients defined according to the WHO 2008 classification. Blood. American Society of Hematology; 2011;117:3163–71.

39. Fry TJ, Shah NN, Orentas RJ, Stetler-Stevenson M, Yuan CM, Ramakrishna S, et al. CD22-targeted CAR T cells induce remission in B-ALL that is naive or resistant to CD19-targeted CAR immunotherapy. Nat Med. 2018;24:20–8.

40. Ruella M, Barrett DM, Kenderian SS, Shestova O, Hofmann TJ, Perazzelli J, et al. Dual CD19 and CD123 targeting prevents antigen-loss relapses after CD19-directed immunotherapies. J Clin Invest. 2016;126:3814–26.

41. Le Magnen C, Shen MM, Abate-Shen C. Lineage Plasticity in Cancer Progression and Treatment. Annu Rev Cancer Biol. 2018;2:271–89.

42. Sadelain M, Brentjens R, Rivière I. The promise and potential pitfalls of chimeric antigen receptors. Current Opinion in Immunology. 2009;21:215–23.

43. Kochenderfer JN, Feldman SA, Zhao Y, Xu H, Black MA, Morgan RA, et al. Construction and Preclinical Evaluation of an Anti-CD19 Chimeric Antigen Receptor. J Immunother. 2009;32:689–702.

44. Zhong X-S, Matsushita M, Plotkin J, Rivière I, Sadelain M. Chimeric antigen receptors combining 4-1BB and CD28 signaling domains augment PI3kinase/AKT/Bcl-XL activation and CD8+ T cell-mediated tumor eradication. Mol Ther. 2010;18:413–20.

45. Gade TPF, Hassen W, Santos E, Gunset G, Saudemont A, Gong MC, et al. Targeted Elimination of Prostate Cancer by Genetically Directed Human T Lymphocytes. Cancer Research. 2005;65:9080–8.

46. Dudley ME, Wunderlich JR, Robbins PF, Yang JC, Hwu P, Schwartzentruber DJ, et al. Cancer regression and autoimmunity in patients after clonal repopulation with antitumor lymphocytes. Science. American Association for the Advancement of Science; 2002;298:850–4.

47. Riddell SR, Greenberg PD. The use of anti-CD3 and anti-CD28 monoclonal antibodies to clone and expand human antigen-specific T cells. Journal of Immunological Methods. 1990;128:189–201.

48. Rockne RC, Hawkins-Daarud A, Swanson KR, Sluka JP, Glazier JA, Macklin P, et al. The 2019 mathematical oncology roadmap. Phys Biol. IOP Publishing; 2019;16:041005–34.

49. Gattinoni L, Zhong X-S, Palmer DC, Ji Y, Hinrichs CS, Yu Z, et al. Wnt signaling arrests effector T cell differentiation and generates CD8+ memory stem cells. Nat Med. 2009;15:808–13.

50. Toor AA, Chesney A, Zweit J, Reed J, Hashmi SK. A dynamical systems perspective on chimeric antigen receptor T-cell dosing. Bone Marrow Transplant. Nature Publishing Group; 2018;371:1507–489.

## References

[1] Kochenderfer JN, Feldman SA, Zhao Y, Xu H, Black MA, Morgan RA, et al. Construction and Preclinical Evaluation of an Anti-CD19 Chimeric Antigen Receptor. J Immunother. 2009; 32:689–702.

[2] Gade TPF, Hassen W, Santos E, Gunset G, Saudemont A, Gong MC, et al. Targeted Elimination of Prostate Cancer by Genetically Directed Human T Lymphocytes. Cancer Research. 2005; 65:9080–8.

[3] Novershtern, N., Subramanian, A., Lawton, L. N., Mak, R. H., Haining, W. N., McConkey, M. E., Habib, N., Yosef, N., Chang, C. Y., Shay, T., Frampton, G. M., Drake, A. C. B., Leskov, I., Nilsson, B., Preffer, F., Dombkowski, D., Evans, J. W., Liefeld, T., Smutko, J. S., Chen, J., Friedman, N., Young, R. A., Golub, T. R., Regev, A., and Ebert, B. L. Densely interconnected transcriptional circuits control cell states in human hematopoiesis. Cell 144(2), 296–309, January (2011).

[4] Ye, Z.-j., Kluger, Y., Lian, Z., and Weissman, S. M. Two types of precursor cells in a multipotential hematopoietic cell line. Proc Natl Acad Sci USA 102(51), 18461–18466, December (2005).

[5] Travers, H., Spotswood, H. T., Moss, P. A. H., and Turner, B. M. Human CD34+ hematopoietic progenitor cells hyperacetylate core histones in response to sodium butyrate, but not trichostatin A. Exp. Cell Res. 280(2), 149–158, November (2002).

[6] Seita, J. and Weissman, I. L. Hematopoietic stem cell: self-renewal versus differentiation. Wiley Interdiscip Rev Syst Biol Med 2(6), 640–653, October (2010).

[7] Fry, T. J., Shah, N. N., Orentas, R. J., Stetler-Stevenson, M., Yuan, C. M., Ramakrishna, S., Wolters, P., Martin, S., Delbrook, C., Yates, B., Shalabi, H., Fountaine, T. J., Shern, J. F., Majzner, R. G., Stroncek, D. F., Sabatino, M., Feng, Y., Dimitrov, D. S., Zhang, L., Nguyen, S., Qin, H., Dropulic, B., Lee, D. W., and Mackall, C. L. CD22-targeted CAR T cells induce remission in B-ALL that is naive or resistant to CD19-targeted CAR immunotherapy. Nat Med 24(1), 20–28, January (2018).

[8] Nitschke, L., Floyd, H., Ferguson, D., and Crocker, P. R. Identification of CD22 ligands on bone marrow sinusoidal endothelium implicated in CD22-dependent homing of recirculating B cells. J. Exp. Med. 189(9), 1513–1518 (1999).

[9] Ruella, M., Barrett, D. M., Kenderian, S. S., Shestova, O., Hofmann, T. J., Perazzelli, J., Klichinsky, M., Aikawa, V., Nazimuddin, F., Kozlowski, M., Scholler, J., Lacey, S. F., Melenhorst, J. J., Morrissette, J. J. D., Christian, D. A., Hunter, C. A., Kalos, M., Porter, D. L., June, C. H., Grupp, S. A., and Gill, S. Dual CD19 and CD123 targeting prevents antigen-loss relapses after CD19-directed immunotherapies. J Clin Invest 126(10), 3814–3826, October (2016).

[10] qiang Ji, X., hua Ji, Z., Shao, H.-J., Wang, H.-Y., He, Y.-X., Zhu, H., and Shao, X.-J. Expression of CD123 in childhood B-lineage actue lymhoblastic leukemia and the application of CD123 in minimal residual disease detection. J Jiangsu Univ. (Medicine Edition) 22(5), 418–421 (2012).

[11] Kong, Y., Huang, X.-J., Hao, L., Qin, Y.-Z., Jiang, Q., Jiang, H., and Liu, Y.-R. CD34^+^CD19^+^ cells with CD123 overexpression are a novel progenostic marker in Ph chromosome-positive acute lymphoblastic leukemia. Zhongguo Shi Yan Xue Ye Xue Za Zhi 22(1), 6–10 (2014).

[12] Velten, L., Haas, S. F., Raffel, S., Blaszkiewicz, S., Islam, S., Hennig, B. P., Hirche, C., Lutz, C., Buss, E. C., Nowak, D., Boch, T., Hofmann, W.-K., Ho, A. D., Huber, W., Trumpp, A., Essers, M. A. G., and Steinmetz, L. M. Human haematopoietic stem cell lineage commitment is a continuous process. Nat Cell Biol 19(4), 271–281, March (2017).

[13] Zhao, X., Gao, S., Wu, Z., Kajigaya, S., Feng, X., Liu, Q., Townsley, D. M., Cooper, J., Chen, J., Keyvanfar, K., Fernandez Ibanez, M. D. P., Wang, X., and Young, N. S. Single-cell RNA-seq reveals a distinct transcriptome signature of aneuploid hematopoietic cells. Blood 130(25), 2762–2773, December (2017).

[14] Ji, H., Ehrlich, L. I. R., Seita, J., Murakami, P., Doi, A., Lindau, P., Lee, H., Aryee, M. J., Irizarry, R. A., Kim, K., Rossi, D. J., Inlay, M. A., Serwold, T., Karsunky, H., Ho, L., Daley, G. Q., Weissman, I. L., and Feinberg, A. P. Comprehensive methylome map of lineage commitment from haematopoietic progenitors. Nature 467(7313), 338–342, September (2010).

[15] Flavahan, W. A., Gaskell, E., and Bernstein, B. E. Epigenetic plasticity and the hallmarks of cancer. Science 357(6348), eaal2380–10, July (2017).

[16] Probst, A. V., Dunleavy, E., and Almouzni, G. Epigenetic inheritance during the cell cycle. Nat Rev Mol Cell Biol 10(3), 192–206, February (2009).

[17] Huang, R. and Lei, J. Dynamics of gene expression with positive feedback to histone modifications at bivalent domains. Int. J. Mod. Phys. B 4, 1850075, November (2017).

[18] Jiao, X. and Lei, J. Dynamics of gene expression based on epigenetic modifications. Communications in Information and Systems 18(3), 125–148 (2018).

